# Ecdysteroid kinase-like (EcKL) paralogs confer developmental tolerance to caffeine in *Drosophila melanogaster*

**DOI:** 10.1101/2021.09.15.460555

**Authors:** Jack L. Scanlan, Paul Battlay, Charles Robin

**Author notes:** Corresponding author: Charles Robin Building 184, School of BioSciences, The University of Melbourne, Parkville Campus, Melbourne, Victoria, 3010, Australia.

## Abstract

A unique aspect of metabolic detoxification in insects compared to other animals is the presence of xenobiotic phosphorylation, about which little is currently understood. Our previous work raised the hypothesis that members of the taxonomically restricted ecdysteroid kinase-like (EcKL) gene family encode the enzymes responsible for xenobiotic phosphorylation in the model insect *Drosophila melanogaster* (Diptera: Ephydroidea)—however, candidate detoxification genes identified in the EcKL family have yet to be functionally validated. Here, we test the hypothesis that EcKL genes in the rapidly evolving Dro5 clade are involved in the detoxification of plant and fungal toxins in *D. melanogaster*. The mining and reanalysis of existing data indicated multiple Dro5 genes are transcriptionally induced by the plant alkaloid caffeine and that adult caffeine susceptibility is associated with a novel naturally occurring indel in *CG31370* (Dro5-8) in the Drosophila Genetic Reference Panel (DGRP). CRISPR-Cas9 mutagenesis of five Dro5 EcKLs substantially decreased developmental tolerance of caffeine, while individual overexpression of two of these genes—*CG31300* (Dro5-1) and *CG13659* (Dro5-7)—in detoxification-related tissues increased developmental tolerance. In addition, we found Dro5 loss-of-function animals also have decreased developmental tolerance of the fungal secondary metabolite kojic acid. Taken together, this work provides the first compelling functional evidence that EcKLs encode detoxification enzymes and suggests that EcKLs in the Dro5 clade are involved in the metabolism of multiple ecologically relevant toxins in *D. melanogaster*. We also propose a biochemical hypothesis for EcKL involvement in caffeine detoxification and highlight the many unknown aspects of caffeine metabolism in *D. melanogaster* and other insects.

**Highlights:**

- Phosphorylation is an under-characterised detoxification reaction in insects
- Dro5 EcKL genes are good detoxification candidate genes in *Drosophila melanogaster*
- Knockout and misexpression of some Dro5 genes modulated tolerance to caffeine
- Dro5 genes may also confer tolerance to the fungal toxin kojic acid
- Caffeine tolerance could be adaptive for *Drosophila* associating with *Citrus* fruits

## 1. Introduction

Metabolic detoxification (also called ‘xenobiotic metabolism’; herein called ‘detoxification’) is the process by which toxic compounds from the environment— often the diet—are chemically modified by an organism such that their toxicity is reduced and/or they can be rapidly excreted from the body (Omiecinski et al., 2011; Williams, 1951). Detoxification is a key aspect of the chemical ecology of insects, where it often defines a species’ niche through an attenuation of the fitness effects of toxins present in food sources (Ibanez et al., 2012) or produced by competitors (LeBrun et al., 2014; Trienens et al., 2017). In addition, resistance to synthetic insecticides often evolves through novel or enhanced detoxification abilities (Joußen et al., 2012; Schmidt et al., 2017; Zhu et al., 2010), making understanding the biochemistry, physiology and genetics of detoxification in insects crucial for the sustainable control of agricultural pests and vectors of human disease.

Conventionally, detoxification as a biochemical process has been conceptually divided into two or three ‘phases’, each of which involves the action of enzymes or transporter proteins (Omiecinski et al., 2011; Williams, 1959, 1951). Phase I— modification—is the addition of functional groups, or cleavage revealing functional groups, that facilitates the addition of further moieties; modification reactions are frequently catalysed by members of the cytochrome P450 and carboxylcholinesterase families (Bernhardt, 2006; Oakeshott et al., 2005). Phase II— conjugation—is the addition of bulky, typically hydrophilic moieties that decreases toxicity and facilitates excretion; conjugation reactions are frequently catalysed by members of the glutathione S-transferase and UDP-glycosyltransferase families (Bock, 2016; Enayati et al., 2005). Phase III—excretion—involves the efflux of toxins and their metabolites out of target cells and tissues, typically mediated by ABC transporters (Wu et al., 2019). Detoxification is thought to mainly occur in three tissues in the insect body—the midgut, the Malpighian tubules and the fat body (Li et al., 2019; Yang et al., 2007)—partially due to the substantial enrichment of detoxification gene expression and xenobiotic metabolism at these sites.

Despite this knowledge, many aspects of the biochemistry and physiology of detoxification in insects remains under-explored. Notably, many insect taxa can phosphorylate xenobiotic molecules, particularly steroidal, phenolic and glycosidic compounds (reviewed in Scanlan et al., 2020), raising the possibility that phosphorylation is a unique Phase II detoxication reaction in insects, at least with respect to other animals (Mitchell, 2015). However, due to a distinct lack of focus on these reactions in the published literature, the toxicological importance of xenobiotic phosphorylation is unclear.

The ecdysteroid kinase-like (EcKL) gene family encodes a group of predicted small-molecule kinases predominantly present in insect and crustacean genomes (Mitchell et al., 2014) that have had limited functional characterisation, with known links between individual genes and ecdysteroid hormone metabolism in the silkworm moth *Bombyx mori* (Lepidoptera: Bombycoidea; Sonobe et al., 2006) and *Wolbachia*-mediated cytoplasmic incompatibility (Liu et al., 2014) in the vinegar fly *Drosophila melanogaster* (Diptera: Ephydroidea). Recently, we proposed that the EcKL family encodes the kinases responsible for xenobiotic phosphorylation in insects, supporting this hypothesis by analysing genomic and transcriptomic data in the genus *Drosophila* (Scanlan et al., 2020). We found that EcKL genes evolve in a rapid birth-death pattern characteristic of other detoxification gene families, are enriched for expression in detoxification-related tissues, and are transcriptionally induced by feeding on xenobiotic compounds; overall, 47% of EcKL genes in *D. melanogaster* have a high ‘detoxification score’, a novel predictive metric validated against the known functions of members of the cytochrome P450 gene family (Scanlan et al., 2020). These data motivated the following experiments to functionally validate the involvement of the EcKL gene family in detoxification processes.

EcKLs in the *Drosophila* genus can be grouped into 46 clades, each derived from a single gene in the most recent common ancestor of the 12 *Drosophila* species we previously examined (Scanlan et al., 2020). These clades (and subclades) give each *Drosophila* EcKL a ‘DroX-Y’ designation (where X is the clade number and Y is the subclade number) for easier comparisons between species, although this is not intended to be used as an official gene name (Scanlan et al., 2020; Jack L. Scanlan, PhD thesis, The University of Melbourne, 2020). One ancestral EcKL clade in *Drosophila*—Dro5—has experienced the largest number of gene duplications (20) in the genus and could be further divided into at least 12 subclades (Fig. 1B). *D. melanogaster* possesses seven genes in the Dro5 clade—*CG31300* (Dro5-1), *CG31104* (Dro5-2), *CG13658* (Dro5-5), *CG11893* (Dro5-6), *CG13659* (Dro5-7), *CG31370* (Dro5-8) and *CG31436* (Dro5-10)—the first five of which are predicted to be involved in detoxification processes based on their detoxification score (Scanlan et al., 2020). These Dro5 paralogs are grouped into two clusters of four and three genes within a larger 26-gene cluster of EcKLs on chromosome 3R (Fig. 1A) and differ in their enrichment within the main detoxification tissues (Fig. 1C), as well as in their induction by the ingestion of xenobiotic compounds or toxic fungal competitors (reviewed in Scanlan et al., 2020). We noticed that three Dro5 genes in *D. melanogaster* are consistently transcriptionally induced in 3^rd^-instar larvae after feeding on the insecticidal plant alkaloid caffeine in two independent datasets (Fig. 1D), raising the possibility that some of these genes may be involved in caffeine metabolism.

**Figure 1.**
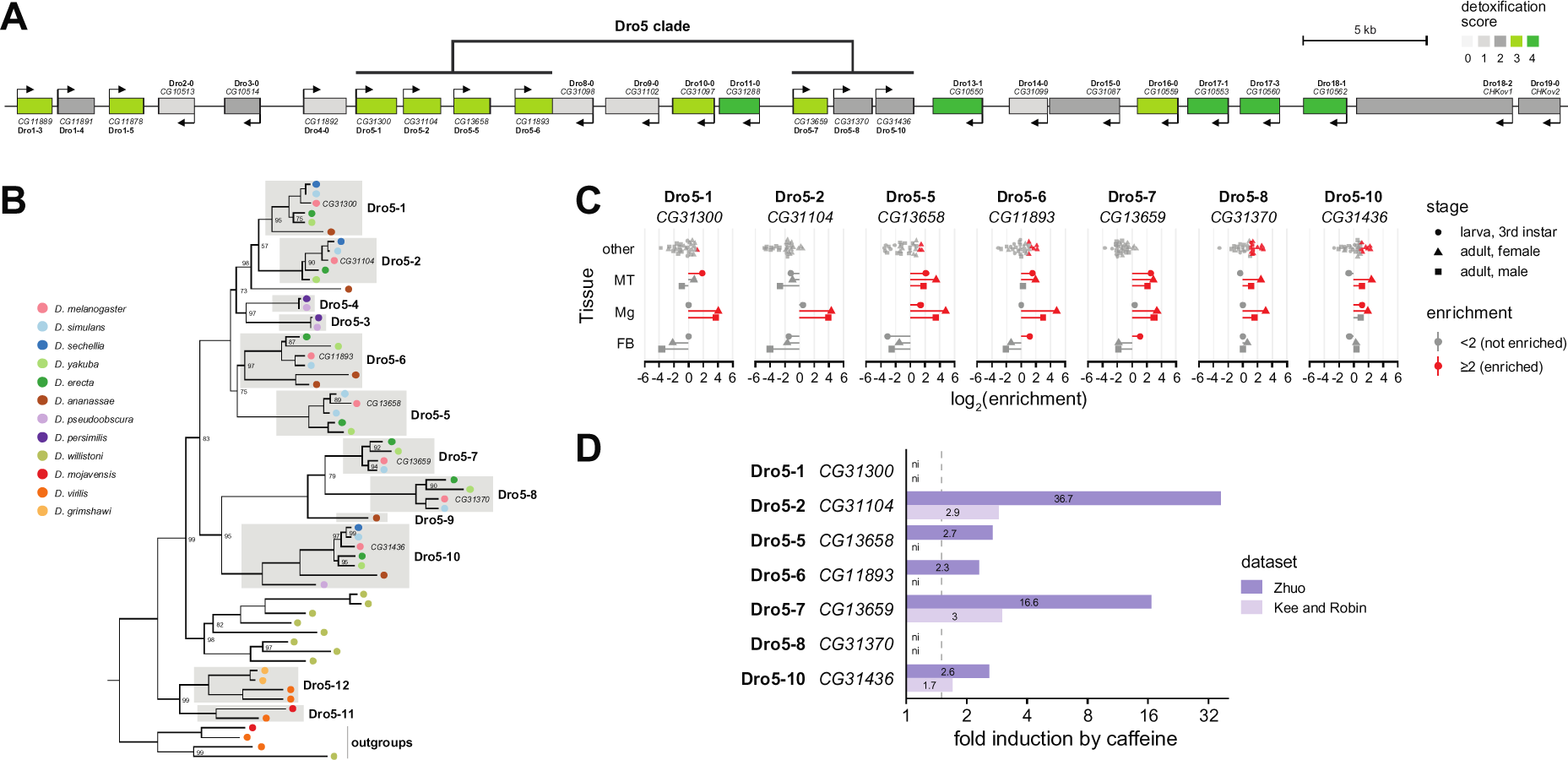
Genomic location, evolution and expression of the Dro5 clade in *Drosophila melanogaster*. (A) The large EcKL gene cluster on chromosome 3R, with the seven Dro5 clade genes indicated. Genes are coloured by their ‘detoxification score’, where a value of 3 or 4 suggests the gene encodes a detoxification enzyme (Scanlan et al., 2020). (B) Phylogenetic tree of 57 Dro5 EcKLs (and four outgroup EcKLs) from 12 *Drosophila* species, grouped into sub-clades (Scanlan et al., 2020). Numbers at nodes are bootstrap support values from UFBoot2 (Hoang et al., 2018); nodes without numbers have support values of 100. Only Dro5 genes from *D. melanogaster* have been named for ease of interpretation, but tips are coloured by the species of origin. (C) Tissue expression enrichment (for detail, see Scanlan et al., 2020) of *D. melanogaster* Dro5 genes in detoxification tissues (MT, Malpighian tubules; Mg, midgut; FB, fat body) and all other tissues (other) at three life stages, based on data in FlyAtlas 2 (Leader et al., 2018). For a given tissue, enrichment values greater than or equal to 2 (red) indicate a gene is nominally ‘enrichment’, while enrichment values less than 2 (grey) indicate a gene is ‘not enriched’, compared to whole-body expression. (D) Transcriptional induction of Dro5 genes in *D. melanogaster* 3^rd^-instar larvae after feeding on caffeine-supplemented media compared to control media, in the Zhuo dataset (Ran Zhuo, PhD thesis, University of Alberta, 2014; dark purple) and the Robin & Kee (2021) dataset (light purple). The fold induction cutoff (1.5x) is indicated with a dashed line; the fold induction is indicated on each bar. Note the log_2_ scale on the x-axis. ni, not induced.

In this study, we provide the first compelling functional evidence that members of the EcKL gene family encode xenobiotic kinases, by testing the hypothesis that Dro5 EcKLs in *D. melanogaster* are involved in the detoxification of plant and fungal secondary metabolites. Using gene disruption and transgenic overexpression techniques, we show that multiple Dro5 genes confer developmental tolerance to caffeine, and also find an association between a naturally occurring deletion allele and adult susceptibility to caffeine in an inbred panel of genotypes (the DGRP). Additionally, we find that Dro5 genes may confer tolerance to the fungal metabolite kojic acid. These results support the hypothesis that EcKLs encode xenobiotic kinases and suggest that the biochemistry of caffeine metabolism in *D. melanogaster* should be revisited in greater detail.

## 2. Material and methods

### 2.1. Fly genotypes and husbandry

#### 2.1.1. Fly husbandry

For routine stock maintenance, flies were kept on yeast-cornmeal-molasses media (‘standard media’; http://bdsc.indiana.edu/information/recipes/molassesfood.html) at 18 °C, 21 °C or 25 °C in plastic vials sealed with cotton stoppers. All bioassays were conducted at 25 °C. Bioassays that were analysed together (each represented by a different figure or sub-figure in the results) were conducted as a group on the same batch of media at the same time to minimise intra-experiment batch effects.

#### 2.1.2. Fly genotypes

The following fly lines were obtained from the Bloomington Drosophila Stock Center (BDSC): *CG31300^MB00063^* (BL22688), *CG13658^MI03110^* (BL37335), *CG11893^MB00360^* (BL22775), *CG31370^MI07438^* (BL44188), *CG31436^MI01111^* (BL33107), *w^1118^*; *Df(3R)BSC852*/TM6C, *Sb^1^*, *cu^1^* (BL27923), *w\**;; *Sb^1^*/TM3, *actGFP*, *Ser^1^* (BL4534), *hsFLP*, *y^1^*, *w^1118^*;; *nos-GAL4*, UAS-*Cas9* (BL54593), and *y^1^*, *v^1^*, *P{y^+t7.7^=nos-phiC31\int.NLS}X*; *P{y^+t7.7^=CaryP}attP40*; (BL25709). DGRP lines were also obtained from the BDSC. The *w^1118^*; *Kr^IF-1^*/CyO; *Sb^1^*/TM6B, *Antp^Hu^*, *Tb^1^* double-balancer line (also known as *w^1118^*-DB), *6g1HR-6c-GAL4* (also known as HR-GAL4; Chung et al., 2007) and *tub-GAL4*/TM3, *actGFP*, *Ser^1^* were a kind gift of Philip Batterham (The University of Melbourne) and Trent Perry (The University of Melbourne). *Df(3R)BSC852*/TM3, *actGFP*, *Ser^1^* was made by crossing BL7923 to BL4534 and selecting the appropriate genotype. *w^1118^*; *Kr^If-1^*/CyO; *nos-GAL4*, UAS-*Cas9* was made by routine crosses, starting with BL54593 males and *w^1118^*-DB females, until the desired genotype was achieved. *w^1118^*; 25709; *Sb^1^*/TM6B, *Antp^Hu^*, *Tb^1^* (chromosome 2 isogenic to BL25709) was made by routine crosses, starting with BL25709 males and *w^1118^*-DB females, until the desired genotype was achieved. *w^1118^*; 25709; *Sb^1^*/TM3, *actGFP*, *Ser^1^* was made by routine crosses, starting with BL25709 males and *w^1118^*; *Kr^IF-1^*/CyO; *Sb^1^*/TM3, *actGFP*, *Ser^1^* females (which themselves were made by routine crosses beginning with BL4534 males and *w^1118^*-DB females), until the desired genotype was achieved.

### 2.2. Plasmid cloning and D. melanogaster transgenesis

#### 2.2.1. pCFD6 cloning

20 nt gRNAs were designed with CRISPR Optimal Target Finder (Gratz et al., 2014; http://targetfinder.flycrispr.neuro.brown.edu/) with the stringency set to ‘maximum’. The pCFD6 vector (Addgene plasmid #73915; http://n2t.net/addgene:73915; RRID:Addgene_73915) was a gift from Simon Bullock. The recombinant pCFD6 plasmids ‘pCFD6-Dro5A’ and ‘pCFD6-Dro5B’, each of which express—under the control of a UAS promoter—four gRNAs that target either the Dro5A or Dro5B locus (Fig. 2A), were designed *in silico* using Benchling (http://benchling.com), and cloned as described by Port & Bullock (2016), with minor modifications below. pCFD6 was digested with *Bbs*I-HF (NEB) and the 9.4 kb backbone gel-purified. The intact pCFD6 vector was used as a template for the production of the three overlapping gRNA-containing inserts by PCR with Phusion Flash polymerase (NEB) using pairs of primers (pCFD6-Dro5A: pCFD6_D5ΔA_1F/R, pCFD6_D5ΔA_2F/R and pCFD6_D5ΔA_3F/R; pCFD6-Dro5B: pCFD6_D5ΔB_1F/R, pCFD6_D5ΔB_2F/R and pCFD6_D5ΔB_3F/R). Gel-purified inserts were cloned into the digested pCFD6 backbone using Gibson assembly (E5520S, NEB) with a 3:1 molar ratio of each insert to vector and 0.3 pmol of total DNA per reaction, with a 4 hr incubation time. 2 μL of each 20 μL assembly reaction was used to transform DH5-*E. coli* (C2987H, NEB), resultant colonies of which were screened for successful assembly with colony PCR —2 min initial denaturation (95 °C), then 2 min denaturation (95 °C), 45 sec annealing (58 °C) and 1 min extension (72 °C) for 32 cycles, then a 5 min final extension (72 °C)—using GoTaq Green Master Mix (M7123, Promega) and the pCFD6_seqfwd and pCFD6_seqrev primers (Table S1) with an expected amplicon size of 890 bp for both plasmids. Plasmids from positive colonies were Sanger sequenced using the pCFD6_seqfwd and pCFD6_seqrev primers at the Australian Genome Research Facility (AGRF).

**Figure 2.**
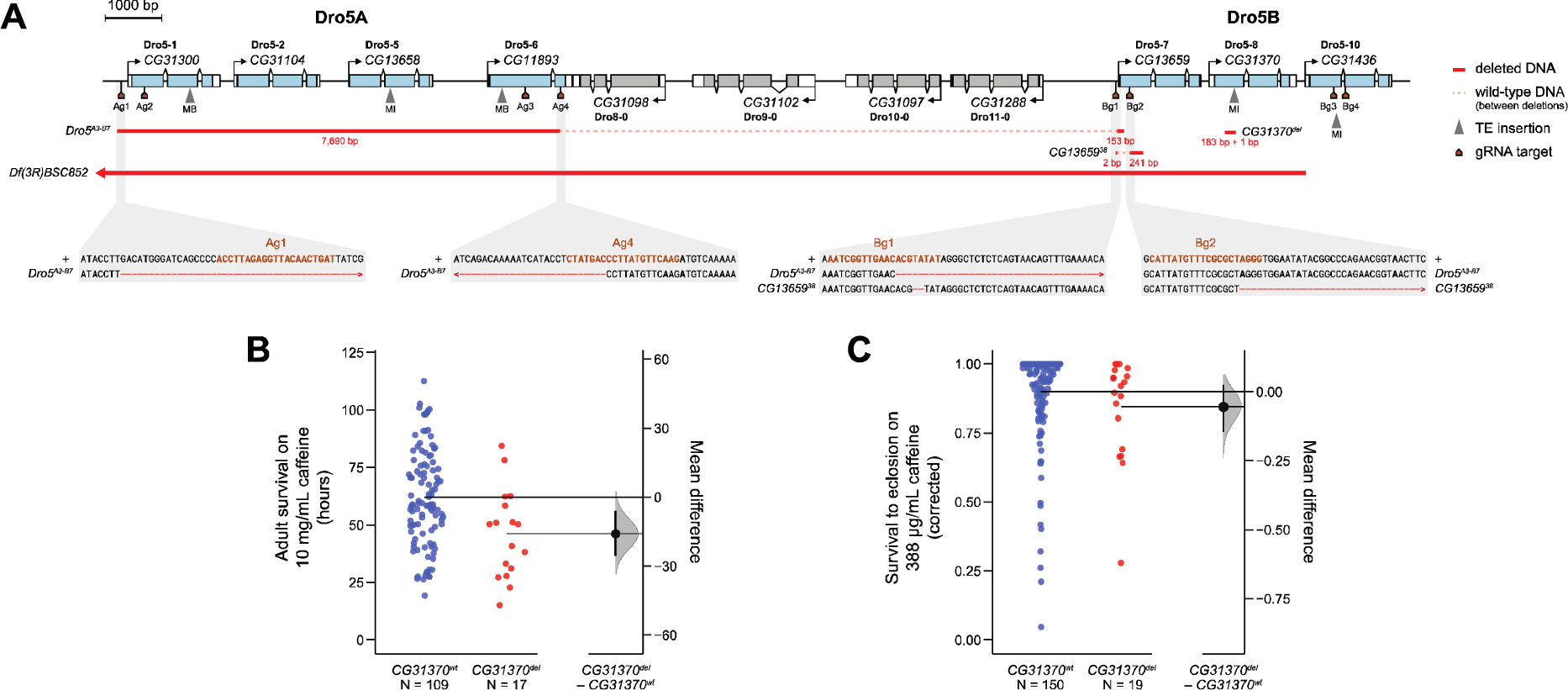
Alleles at the Dro5A and Dro5B loci in *D. melanogaster* and caffeine-related phenotypes of *CG31370* genotypes in the DGRP. (A) The location of deletion and transposable element (TE) insertion alleles in Dro5 genes (blue) at the Dro5A and Dro5B loci, either induced by CRISPR-Cas9 mutagenesis in this study (*Dro5^A3-B7^* and *CG13659^38^*), previously derived from TE insertion (MI and MB) or FRT-mediated deletion (*Df(3R)BSC852*), or naturally present in the DGRP (*CG31370^del^*). Top: Gene models, with coding sequence in blue (Dro5 genes) or grey (other EcKLs) and non-coding (UTR) sequence in white. Middle: Positions and sizes of molecular lesions. Bottom: Sequence-level detail of deleted nucleotides in CRISPR-Cas9-derived alleles compared to the wild-type genetic background (+), with the gDNA target sites highlighted in brown. (B,C) Estimation plots (Ho et al., 2019) of mean phenotypic differences between homozygous *CG31370^wt^* (blue) and homozygous *CG31370^del^* (red) DGRP lines. The right-hand axis shows the mean difference (effect size; black dot) between groups, with the 95% CI (black line) and distribution of bootstrapped means (grey curve). Effect sizes with CIs that do not include zero are considered significant. (B) Survival in hours of adult female flies of different DGRP lines on 10 mg/mL caffeine-supplemented media (as phenotyped by Najarro et al., 2015). (C) Corrected proportional survival to eclosion of larvae of different DGRP lines developing in 388 µg/mL caffeine-supplemented media (as phenotyped by Montgomery et al., 2014).

**Table 1.** Collated evidence for the involvement of individual *Drosophila melanogaster* Dro5 EcKLs in caffeine tolerance.

| Gene | Nomenclature | Induction (Z)<sup>a</sup> | Induction (K)<sup>b</sup> | DGRP<sup>c</sup> | *Dro5*<sup>A3-B7</sup><sup>d</sup> | KO L-A<sup>e</sup> | KO L-P<sup>f</sup> | KO P-A<sup>g</sup> | Misexpression<sup>h</sup> |
| --- | --- | --- | --- | --- | --- | --- | --- | --- | --- |
| CG31300 | Dro5-1 | No | No | - | Yes | Yes | No | Yes | Yes |
| CG31104 | Dro5-2 | Strong | Weak | - | Yes | N.D. | N.D. | N.D. | No |
| CG13658 | Dro5-5 | Weak | No | - | Yes | Yes | No | Yes | No |
| CG11893 | Dro5-6 | Weak | No | - | Yes | Yes | Yes | Yes | N.D |
| CG13659 | Dro5-7 | Strong | Weak | - | Yes | Yes | Yes | Yes | Yes |
| CG31370 | Dro5-8 | No | No | Yes | - | N.D | N.D. | N.D. | No |
| CG31436 | Dro5-10 | Weak | Weak | - | - | N.D. | N.D. | N.D. | N.D |

<sup>a</sup> Induction by 1,553 µg/mL caffeine in the Zhuo dataset (Fig. 1D); strong, > 3-fold; weak, ≤ 3-fold
<sup>b</sup> Induction by 1,500 µg/mL caffeine in Robin & Kee dataset (Fig. 1D); strong, > 3-fold; weak, ≤ 3- fold
<sup>c</sup> Association of genetic variation with caffeine-related phenotypes in the DGRP (Fig. 2); -, no associated variation
<sup>d</sup> Disrupted in the *Dro5*<sup>A3-B7</sup> allele, which is associated with increased developmental susceptibility to caffeine (Figs. 2 & 4); -, not disrupted
<sup>e</sup> Single-gene knockout (KO) increases larval–adult developmental caffeine susceptibility (Figs. 5A & S3); N.D., not determined
<sup>f</sup> Single-gene knockout (KO) increases larval–pupal developmental caffeine susceptibility (Figs. 5A & S3); N.D., not determined
<sup>g</sup> Single-gene knockout (KO) increases pupal–adult developmental caffeine susceptibility (Figs. 5A & S3); N.D., not determined
<sup>h</sup> Transgenic misexpression in detoxification tissues increases developmental caffeine tolerance (Fig. 5B & S4); N.D., not determined

#### 2.2.2. pUASTattB cloning

Full-length cDNA clones for *CG31300* (FI01822), *CG31104* (IP12282), *CG13658* (FI12013), *CG11893* (IP11926), *CG13659* (IP11858), *CG31370* (IP10876) and *CG31436* (IP12392) were obtained from the Drosophila Genomics Resource Center (DGRC). Recombinant pUASTattB plasmids (Bischof et al., 2007) containing individual EcKL ORFs under the control of a UAS promoter were designed *in silico* using Benchling (http://benchling.com). The pUASTattB vector was digested with *Eag*I-HF (NEB) and *Kpn*I-HF (NEB) and the 8.5 kb backbone gel-purified (28704, Qiagen). ORFs were amplified with PCR—10 sec initial denaturation (98 °C), then 5 sec denaturation (98 °C), 5 sec annealing (55 °C) and 15 sec extension (72 °C) for 32 cycles, then a final 1 min extension (72 °C)—using Phusion Flash polymerase (NEB) from cDNA clones using primers containing an *Eag*I restriction site (forward primers) or a *Kpn*I restriction site (reverse primers), as well as an additional 5’ sequence (5’-TAAGCA-3’) to aid digestion (Table S1). Amplicons were column-purified (FAPCK 001, Favorgen), double-digested with *Eag*I-HF and *Kpn*I-HF for 8 hr, then gel-purified. *Eag*I/*Kpn*I-digested ORFs were ligated into the *Eag*I/*Kpn*I-digested pUASTattB vector backbone using a 6:1 insert:vector molar ratio and T4 DNA ligase (M0202S, NEB) in a thermocycler overnight (∼16 hr), alternating between 10 °C for 30 sec and 30 °C for 30 sec (Lund et al., 1996). 5 μL of each 20 μL ligation reaction was used to transform DH5-*E. coli*, resultant colonies of which were screened for successful assembly with colony PCR—2 min initial denaturation (95 °C), then 2 min denaturation (95 °C), 45 sec annealing (58 °C) and 1.5 min extension (72 °C) for 32 cycles, then a 5 min final extension (72 °C)—using GoTaq Green Master Mix and the pUASTattB_3F/5R primers (Table S1). Plasmids from positive colonies were Sanger sequenced using the pUASTattB_3F and pUASTattB_5R primers at AGRF.

#### 2.2.3. *D. melanogaster* transgenesis

Correctly assembled plasmids were sent to TheBestGene Inc. (US) for microinjection and incorporation into the *D. melanogaster* genome at the attP40 site on chromosome 2 (BL25709). Transformed lines were received as a mixture of white-eyed (zero copies of the mini-*white* gene), orange-eyed (one copy of the mini-*white* gene) and red-eyed (two copies of the mini-*white* gene) flies—virgin white-eyed flies were pooled and retained as a genetic background line (‘yw’), while the plasmid-integrated lines were individually kept as the red-eyed homozygous stocks ‘pCFD6Dro5A’ and ‘pCFD6Dro5B’ (of genotype *w^-^*, 25709; *pCFD6*; 25709), and UAS-*CG31300*, UAS-*CG31104*, UAS-*CG13658*, UAS-*CG11893*, UAS-*CG13659*, UAS-*CG31370* and UAS-*CG31436* (of genotype *w^-^*, 25709; *pUASTattB*; 25709).

### 2.3. CRISPR-Cas9 mutagenesis

Transgenic CRISPR-Cas9 mutagenesis of wild-type chromosomes was performed with the crossing scheme in Figure S1A. Single founder male flies—heterozygous for possibly mutagenised loci on chromosome 3—were allowed to mate with *w^1118^*; 25709; *Sb^1^*/TM3, *actGFP*, *Ser^1^* virgin females, and when larvae were observed in the food media, the DNA from each founder male was extracted as per Bischof *et al*. (2014) and genotyped by PCR as below. Mutagenesis of already-mutagenised chromosomes was performed with the crossing scheme in Figure S1B, using homozygous mutant lines generated previously. Single founder male flies—which were heterozygous for possibly (singly- or doubly-) mutagenised loci on chromosome 3—were allowed to mate with *w^1118^*; 25709; *Sb^1^*/TM3, *actGFP*, *Ser^1^* virgin females, and when larvae were observed in the food media, the DNA from each founder male was extracted using the squish prep protocol, then PCR genotyped with four GoTaq Green reactions per line, which were combined before gel-purification to allow for the detection of early-cycle polymerase-derived errors by close inspection of the sequencing chromatogram output. Dro5A genotyping used the primer pairs D5ΔA_1F/1R and D5ΔA_2F/2R, and Dro5B genotyping used the D5ΔB_1F/1R and D5ΔB_2F/2R primer pairs. PCR—2 min initial denaturation (95 °C), then 2 min denaturation (95 °C), 45 sec annealing (55 °C) and 1.5 min extension (72 °C) for 32 cycles, then a 5 min final extension (72 °C)—was carried out with GoTaq Green Master Mix. Gel-purified amplicons were sequenced using the appropriate genotyping primers at AGRF.

Wild-type flies with the genetic background of flies bearing Dro5 mutations (*w^1118^*; 25709; 25709—otherwise known as ‘+’) were generated by following the crossing scheme in Figure S1A, but using BL25709 as the maternal genotype in C1 instead of a *pCFD6*-containing line.

### 2.4. Standard media developmental viability assays

#### 2.4.1. Egg-to-adult developmental viability assays

Egg-to-adult viability was estimated from the adult genotypic ratios of successfully eclosing offspring produced from crosses between a homozygous parental genotype and a heterozygous parental genotype, the latter of which had at least one phenotypic marker that revealed the genotype of the offspring. Males and females of the relevant genotypes were allowed to mate and lay eggs on vials of standard media, with at least five vials per cross, and the number of adults of each genotype were scored after development at 25 °C for 14 days. If the adult genotypic ratio was significantly different from the 1:1 Mendelian expectation, as determined by the ‘binom.test’ function in R, this was considered evidence that one genotype was less viable than the other.

#### 2.4.2. Larval-to-adult developmental viability assays

Larval-to-adult viability was estimated by transferring particular quantities of 1st-instar larvae of known genotypes (either as the offspring of a cross between homozygous parents, or offspring sorted phenotypically by the presence or absence of a dominant marker such as GFP expression) to vial of standard media, letting them develop at 25 °C for 14 days, and scoring the number of individuals that reached the stages of pupariation, pupation, pharate adult and eclosion. Fisher’s exact test (‘fisher.test’ function in R) was used to determine if there were significant differences between genotypes.

### 2.5. DGRP analyses

#### 2.5.1. DGRP *in silico* and PCR genotyping

BAM files containing alignments of DGRP line sequences from Illumina platforms to the *y; cn bw sp;* reference genome were recovered from the Baylor College of Medicine website (https://www.hgsc.bcm.edu/content/dgrp-lines; Mackay et al., 2012). Local alignments were visualized with IGV (Thorvaldsdóttir et al., 2013) to manually score structural variation *in silico*. For PCR genotyping, the primers CG31370del_1F and CG31370del_1R (Table S1) were designed to flank the *CG31370^del^* region and produce a 576 bp amplicon from *CG31370^wt^* and a 392 bp amplicon from CG31370^del^. DNA was extracted from single flies as per Bischof *et al*. (2014) with three independent extractions per DGRP line. PCR—2 min initial denaturation (95 °C), then 30 sec denaturation (95 °C), 30 sec annealing (53 °C) and 40 sec extension (72 °C) for 30 cycles, then a 5 min final extension (72 °C)—was carried out with GoTaq Green Master Mix, using 0.4 μL of DNA extract as a template per 10 μL reaction. Amplicons were visualised with gel electrophoresis using 1.5% agarose gels.

#### 2.5.2. DGRP caffeine tolerance data and analyses

Adult caffeine survival data was obtained from Najarro et al. (2015). Developmental caffeine (388 μg/mL) survival data from Montgomery et al. (2014) was averaged across the three replicates, and then corrected for (similarly averaged) control (0 μg/mL caffeine) survival using Abbott’s formula (Abbott, 1925), with corrected survival values greater than 1 (indicating greater survival than control) adjusted to 1. Basal gene expression levels in adult female and adult male flies from different DRGP lines were obtained from Everett et al. (2020). Mean differences in phenotypes and gene expression between *CG31370* genotypes were determined with the *dabestr* package (version 0.2.5; Ho et al., 2019) in R; effect sizes with 95% confidence intervals that did not include zero were considered significant.

### 2.6. Single-dose developmental toxicological assays

#### 2.6.1. Media

Quercetin, escin, esculin and curcumin were purchased from Sigma-Aldrich. Toxin-containing media and control media were prepared by adding 100 µL of quercetin, escin, esculin or curcumin dissolved in 100% EtOH or 100 µL of EtOH, respectively, to 5mL of molten yeast-sucrose media (5% w/v inactive yeast, 5% w/v sucrose, 1% w/v agar, 0.38% v/v propionic acid, 0.039% v/v orthophosphoric acid, 0.174% w/v Tegosept, 1.65% v/v EtOH) in each vial and mixing with a clean plastic rod. Media was stored at 4 °C for a maximum of three days before use.

#### 2.6.2. Assays

*Dro5^A3-B7^* females were mated to *Dro5^A3-B7^* or wild-type (+; the genetic background of the CRISPR-Cas9 mutagenesis lines) males and were allowed to lay on apple juice plates (2% w/v agar, 3.125% w/v sucrose, 25% v/v apple juice) topped with yeast paste. After hatching, 20 1st-instar larvae were transferred to each vial of media using a fine paintbrush that was washed between each transfer, and left to develop at 25 °C for 14 days. Vials were scored for the number of individuals that had pupated (formation of the puparium) and that had successfully eclosed (complete vacation of the puparium). Mortality counts were determined as ‘larval’ (# of larvae – # of pupae) or ‘pre-adult’ (# of larvae – # of adults eclosed), and proportional mortality was calculated by dividing mortality counts by the number of larvae added to each vial. Mean differences in proportional mortality between the two genotypes on each type of media were analysed with Welch’s two-sided t-test with unequal variance (‘t.test’ function in R).

### 2.7. Multiple-dose developmental toxicological assays

#### 2.7.1. Media

Caffeine, kojic acid and salicin were purchased from Sigma-Aldrich. Toxin-containing media was prepared by adding toxin stock solution—compound dissolved in dH_2_O— to molten 1.25x yeast-sucrose media (Section 2.6.1): X μL of 40–50 mg/mL toxin stock solution, 1000–X μL of dH_2_O (where X varied according to the final concentration of toxin) and 4 mL of 1.25x media were added to each vial, for a total media volume of 5 mL, then mixed with a clean plastic rod. Control media was made by mixing 4 mL of molten 1.25x media and 1 mL of dH_2_O. Media was stored at 4 °C for a maximum of three days.

#### 2.7.2. Assays

Males and females of the relevant genotypes were crossed, and females were allowed to lay on apple juice plates (Section 2.6.2) topped with yeast paste. 20–30 1st-instar larvae were transferred to each toxin-containing food vial using a fine paintbrush that was washed between each transfer, and left to develop at 25 °C for 14 days. Crosses involving GFP-marked balancer chromosomes had larvae sorted against GFP under a bright-field fluorescent microscope before transferal. Vials were scored for the number of individuals that had pupated (formation of the puparium) and that had successfully eclosed (complete vacation of the puparium). Three types of survival counts (toxicological endpoints) were calculated: ‘larval-to-pupal’ (L-P; # of larvae – # of pupae), ‘pupal-to-adult’ (P-A; # of pupae – # of adults eclosed) or ‘larval-to-adult’ (L-A; # of larvae – # of adults eclosed). Survival counts were converted to proportional survival by dividing by the number of larvae per vial, except in the case of pupal mortality counts, which were converted by dividing by the number of pupae; vials with zero pupae were excluded from PA models to avoid undefined values. Proportional survival data were analysed with the *drc* package (version 3.0-6; Ritz et al., 2015). 3-parameter log-logistic regression models (with a fixed lower limit of 0) were fit with proportional survival as the response and toxin concentration in µg/mL as the dose; all models were assessed with the ‘noEffect’ function to check for a significant dose-response effect. LC_50_ values and their 95% confidence intervals (95% CIs) were calculated relative to the model’s estimated upper limit (ie. the background mortality of each genotype), using robust standard errors from the *sandwich* package (version 2.5-1; Zeileis, 2006, 2004). Statistical comparison of LC_50_ values was performed with the ‘EDcomp’ function in *drc*, with the 95% CI of the ratio of the LC_50_s excluding 1 being considered statistically significant.

### 2.8. Citrus media developmental viability assays

#### 2.8.1. Media

‘Delite’ mandarin oranges (*Citrus reticulata*), ‘Ruby Blush’ grapefruits (*C. × paradisi*) and navel sweet oranges (*C. × sinensis*) were juiced with a hand juicer, juice was strained to remove large pulp particles, and yeast, agar and dH_2_O were added and heated in a microwave. After cooling to 60 °C, 10% Tegosept in EtOH was added, and 5 mL of media was aliquoted into each vial. Final concentrations of yeast and agar were 5% and 1% w/v, respectively, and 0.174% and 1.65% v/v for Tegosept and EtOH, respectively (5 g yeast, 1 g agar, 20 mL dH_2_O and 80 mL juice for 100 mL of media).

#### 2.8.2. Assays

Thirty 1st-instar larvae were transferred to each fruit media vial using a fine paintbrush and left to develop at 25 °C for 14 days. Vials were each scored for the number of larvae that had pupated (formation of the puparium) and that had successfully eclosed (complete vacation of the puparium) and proportional survival was calculated by dividing by the number of larvae added to each vial. Mean differences in proportional mortality between the two genotypes on each type of media was analysed with Welch’s two-sided t-test with unequal variance (‘t.test’ function in R).

### 2.9. Data handling and presentation

Data cleaning and restructuring was performed in R (version 3.6.1; Team, 2019) using the *tidyverse* collection of packages (version 1.2.1; Wickham et al., 2019). Genomic loci were visualised with GenePalette (www.genepalette.org; Smith et al., 2017). Data were visualised with either the *ggplot2* package (version 3.2.1; Wickham, 2016) or the *dabestr* package (version 0.2.5; Ho et al., 2019) in R, and resulting plots were polished in Adobe Illustrator.

## 3. Results

### 3.1. CRISPR-Cas9 mutagenesis of the Dro5 EcKL clade

We attempted to generate a *D. melanogaster* line that had all seven Dro5 genes specifically deleted or disrupted, using CRISPR-Cas9 mutagenesis—a ‘Dros5-null’ allele. As the Dro5 genes lie in two clusters—Dro5A (containing four genes) and Dro5B (containing three genes; Fig. 2A)—separated by four non-Dro5 EcKLs, we used an approach in which a multi-gRNA-expressing pCFD6 construct (Port and Bullock, 2016) targets one cluster of genes, then the resulting deletion alleles is used as the genetic background for another round of mutagenesis with a separate pCFD6 construct (Fig. S1). Mutagenesis at the Dro5A cluster produced eight large deletion alleles (putatively generated by cuts between distant gRNA target sites) detected through PCR from screening 24 founder males, including the *Dro5^A3^* allele, which contained a 7,690 bp deletion (Fig. 2A). Forty founder males were screened for mutations at the Dro5B cluster using PCR, but no large deletions encompassing the entire Dro5B cluster were detected; given a lack of heteroduplex bands generated after PCR with the D5ΔB_2F/D5ΔB_2R primer pair, the 3^rd^ and 4^th^ gRNAs from the pCFD6-Dro5B construct failed to cut. However, we successfully isolated a frameshifting composite deletion allele of *CG13659* (Dro5-7) designated *CG13659^38^* (consisting of a 2 bp deletion 43 bp upstream of the transcription start site and a 241 bp deletion in the first exon that deleted 81 aa; Fig. 2A). The Dro5A allele *Dro5^A3^* was selected for a second round of mutagenesis to produce additional deletions at the Dro5B cluster—16 founder males were screened, with two putative deletions detected at the *CG13659* locus, but none across the Dro5B cluster as a whole. Thus a fly line, *Dro5^A3-B7^*, was generated that was a homozygous-viable composite allele consisting of two deletions, one 7,690 bp long in Dro5A and one 153 bp long in Dro5B. This bore a complete deletion of *CG31300* (Dro5-1), *CG31104* (Dro5-2) and *CG13658* (Dro5-5), and partial deletions of 1,245 bp (415 aa) of the CDS of *CG11893* (Dro5-6) and the first 85 bp (28 aa) of *CG13659* (Dro5-7), including the transcription and translation start sites of both genes (Fig. 2A).

### 3.2. Dro5 EcKLs are not required for normal development of Drosophila melanogaster

To test if any Dro5 genes are required for gross development in *D. melanogaster* on standard media, we placed loss-of-function alleles for all Dro5 genes—either pre-existing transposable element coding sequence (CDS) insertions or those generated in Section 3.1—in *trans* to the homozygous-lethal chromosomal deficiency *Df(3R)BSC852*, which deletes or otherwise likely disrupts all seven Dro5 genes (and 11 other genes), and measured egg-to-adult viability. Loss of five individual Dro5 genes, or five genes simultaneously in the case of the *Dro5^A3-B7^* allele, did not significantly affect adult genotypic ratios, suggesting Dro5 genes are not required for normal development in the absence of toxic challenge (Fig. 3A). A larval-to-adult viability experiment involving just the *Dro5^A3-B7^* allele further supported this conclusion, with the vast majority of *Dro5^A3-B7^*/*Df(3R)BSC852* individuals successfully completing development, and no significant difference was found between the developmental outcomes of *Dro5^A3-B7^*/*Df(3R)BSC852* and *Dro5^A3- B7^*/TM3, *actGFP*, *Ser^1^* animals (p = 0.56, Fisher’s exact test; Fig. 3B).

**Figure 3.**
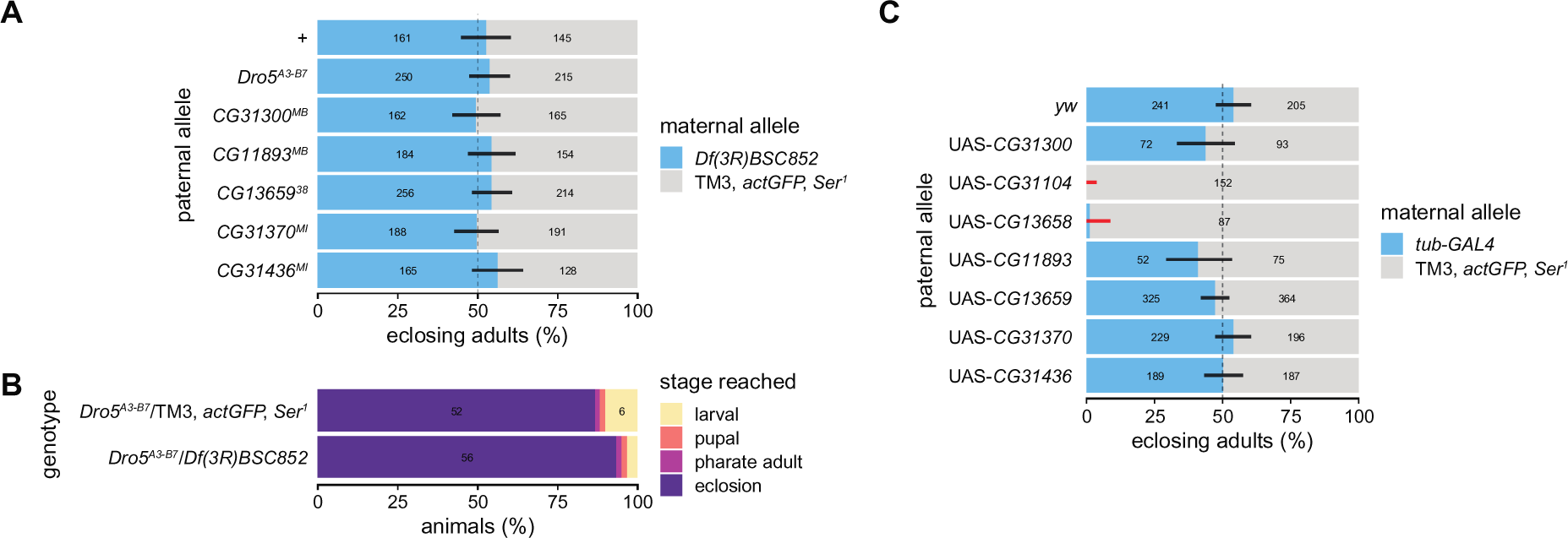
Developmental viability of Dro5 mutants and Dro5 ubiquitous overexpression animals on lab media. (A) Egg-to-adult viability of Dro5 loss-of-function alleles (or the wild-type allele +) over the deficiency *Df(3R)BSC852* estimated from the adult genotypic ratios of offspring from crosses between *Df(3R)BSC852*/TM3, *actGFP*, *Ser^1^* females and males of one of seven homozygous null-allele genotypes. The dashed line indicates the expected 1:1 genotypic ratio if both genotypes per cross are equally developmentally viable; error bars are 95% confidence intervals (adjusted for seven tests) for the proportion of *Df(3R)BSC852* heterozygotes; black and red bars indicate non-significant or significant deviations, respectively, from expected genotypic ratios after correction for multiple tests. Numbers on the bars are the number of adults scored of that genotype. (B) Larvae-to-adult viability of offspring from the cross between *Df(3R)BSC852*/TM3, *actGFP*, *Ser^1^* females and homozygous *Dro5^A3-B7^* males, sorted at the 1st-instar larval stage by GFP fluorescence (n = 60 larvae per genotype). Numbers on the bars are the number of individuals in each lethal phase category (for numbers greater than five). (C) Egg-to-adult viability of the misexpression of Dro5 ORFs (or non-misexpression from the genetic background *yw*) using the strong, ubiquitous GAL4 driver *tub-GAL4*, estimated from the adult genotypic ratios of offspring from crosses between *tub-GAL4*/TM3, *actGFP*, *Ser^1^* females and males of one of eight homozygous responder genotypes. The dashed line indicates the expected 1:1 genotypic ratio if both genotypes per cross are equally developmentally viable; error bars are 99.38% CIs (95% CI adjusted for eight tests) for the proportion of *tub-GAL4* heterozygotes; black and red bars indicate non-significant or significant deviations, respectively, from expected genotypic ratios after correction for multiple tests. Numbers on the bars are the number of adults scored of that genotype.

In addition, all seven Dro5 genes were individually misexpressed from the pUASTattB vector with the strong, ubiquitous GAL4 driver *tub-GAL4* on standard media to test if misexpression could disrupt developmental progression. Misexpression of *CG31104* (Dro5-2) and *CG13658* (Dro5-5) resulted in no or very few successfully eclosing adults (Fig. 3C), suggesting ectopic or excessive expression of either gene arrests development. Examination of *tub*>*CG31104* and *tub*>*CG13658* animals, using brightfield fluorescence microscopy to select against GFP-positive individuals, revealed that these genotypes are arrested during metamorphosis, with pharate adults having completely undifferentiated abdomens, lacking bristles and genitalia. Misexpression of the other five Dro5 genes did not significantly change adult genotypic ratios (Fig. 3C) and therefore does not appear to grossly affect developmental progression.

### 3.3. A composite deletion in the Dro5B cluster further motivates an exploration of caffeine tolerance in D. melanogaster

A manual reanalysis of structural variation associated with EcKL genes in the Drosophila Genetic Reference Panel (DGRP; Mackay et al., 2012) identified a novel composite deletion in the first exon of *CG31370* (Dro5-8) compared to the Release 6 reference genome, of sizes 183 bp (3R:25,302,734..25,302,916) and 1 bp (3R:25,302,921), 5 bp apart. Given that this naturally occurring allele was composed of derived deletions (based on comparisons with CG31370 orthologs in other Drosophila genomes; Scanlan et al., 2020), it was designated *CG31370^del^* (Fig. 2A), while the ancestral allele was designated *CG31370^wt^*. *CG31370^del^* is missing 61 aa of the encoded CG31370 protein and also induces a frameshift, making it a likely strong loss-of-function allele. 202 DGRP lines were genotyped *in silico* at *CG31370*, with 182 lines homozygous for *CG31370^wt^*, 17 lines homozygous for *CG31370^del^* and two lines heterozygous for both alleles; five lines were unable to be called due to uninformative read mapping depth or a lack of available mapped-read data. We also genotyped the *CG31370* locus of 46 DGRP lines using PCR, revealing a single additional *CG31370^wt^* homozygote and four additional *CG31370^del^* homozygotes among the five uncalled lines, as well as validating the *in silico* genotypes of 31 and 10 *CG31370^wt^* homozygotes and *CG31370^del^* homozygotes, respectively. (For the genotypes of all lines, see Table S1.)

We tested whether there was an association between this recharacterized naturally occurring *CG31370^del^* allele and the caffeine susceptibility data generated by Najarro *et al*. (2015). They measured the mean lifespan of adult female flies from 165 DGRP lines feeding on media containing 1% (10 mg/mL) caffeine but found no significant genome-wide associations in their analyses using SNPs; of 165 lines with a phenotype in their dataset, 126 had a confident *CG31370* genotype (109 lines homozygous for *CG31370^wt^*, 17 lines homozygous for *CG31370^del^*, and one heterozygous line). We added the *CG31370^del^* genotype to the DGRP variant data (http://dgrp2.gnets.ncsu.edu/data.html) and performed a GWAS with PLINK (Purcell et al., 2007) using the five major inversion and *Wolbachia* infection status as covariates and the phenotypic data from Najarro et al. (Najarro et al., 2015), and the *CG31370^del^* allele ranked among the top 0.3% of annotated variants in the DGRP. An estimation statistics approach suggests the mean difference in the survival time on 10 mg/mL caffeine between homozygous *CG31370^wt^* and *CG31370^del^* lines was 15.9 hours (95% CI: 5.93, 25.5; Fig. 2B), suggesting the *CG31370^del^* allele increases adult susceptibility to caffeine.

We also tested whether the *CG31370^del^* allele is negatively associated with a developmental caffeine survival phenotype, measured by Montgomery et al. (2014) as successful development (larval feeding through to adult eclosion) on media containing 388 μg/mL caffeine; of the 173 DGRP lines with a phenotype in their dataset, 169 had a confident *CG31370* genotype (150 homozygous for *CG31370^wt^*, 19 homozygous for *CG31370^del^*, and two heterozygous lines). The mean difference in the corrected survival proportion on 388 μg/mL caffeine between homozygous *CG31370^wt^* and *CG31370^del^* lines was 0.054 (95% CI: -0.025, 0.15; Fig. 2C), suggesting these genotypes do not significantly differ in their susceptibility to caffeine at this dose. Given that most DGRP lines showed high corrected survival on 388 µg/mL caffeine, it is likely that this dose—which was originally intended by Montgomery et al. (2014) to be sub-lethal and was 25.8-fold lower than that used by Najarro et al. (2015)—was insufficiently high to discriminate between the *CG31370^wt^* and *CG31370^del^* genotypes, if they do indeed vary in their developmental susceptibility to caffeine.

### 3.4. Dro5 loss-of-function mutants have increased developmental susceptibility to caffeine

To further test the hypothesis that some Dro5 genes function in caffeine detoxification, we conducted dose-response developmental toxicology assays on 50–1,500 µg/mL caffeine media with wild-type and *Dro5^A3-B7^* homozygote animals, with three toxicological endpoints determined: larval-to-adult (L-A) survival, larval-to-pupal (L-P) survival, and pupal-to-adult (P-A) survival. The median lethal concentration (LC_50_) of caffeine was significantly lower for *Dro5^A3-B7^* homozygotes than for wild-type animals, with LC_50_ ratios of 0.26 (95% CI: 0.247, 0.274), 0.656 (95% CI: 0.565, 0.747) and 0.184 (95% CI: 0.158, 0.211) for L-A, L-P and P-A survival, respectively (Fig. 4), indicating increased developmental susceptibility to caffeine in mutant animals, particularly during metamorphosis. A notable, qualitative effect of caffeine exposure on *Dro5^A3-B7^* mutant animals was high pharate adult lethality—where pharate adults attempted to eclose but remained trapped in the puparium before dying—even at relatively low concentrations (200–300 µg/mL) where survival from the larval stage to pupation was high.

**Figure 4.**
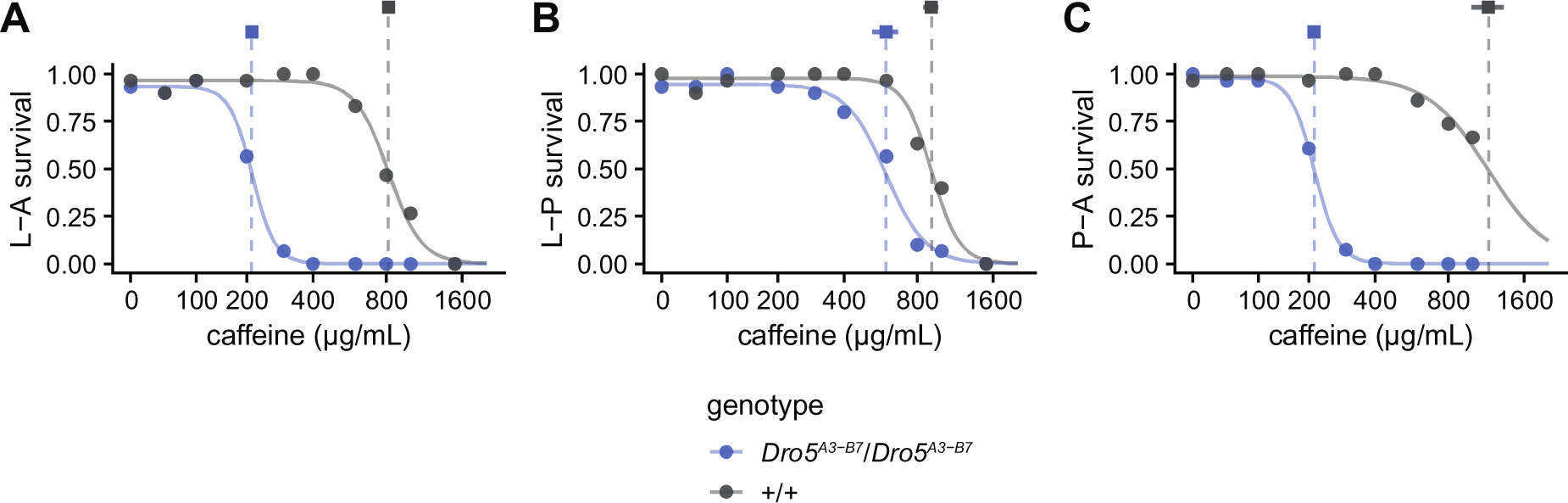
Developmental survival of +/+ homozygotes (grey) and *Dro5^A3-B7^*/*Dro5^A3-B7^* homozygotes (blue) on media containing 0–1,500 µg/mL caffeine. Curves are fitted log-logistic regression dose-response models for each genotype. Dashed vertical lines and squares indicate the estimated LC_50_s for each genotype, with the horizontal bar indicating the 95% CI. (A) L-A (larval-adult) survival. (B) L-P (larval-pupal) survival. (C) P-A (pupal-adult) survival. Curves are fitted log-logistic regression dose-response models for each genotype.

We also performed dose-response developmental toxicology assays with wild-type and *Dro5^A3-B7^* chromosomes in *trans* with the *Df(3R)BSC852* deficiency, which should fully fail to complement the *Dro5^A3-B7^* allele, to confirm that the susceptibility to caffeine seen in *Dro5^A3-B7^* homozygotes was due to mutations at the Dro5 locus and not a secondary site mutation on chromosome 3. There was no significant dose effect for *Df(3R)BSC852*/+ animals at any of the three endpoints for the concentration range used, but there was a large reduction in in L-A and P-A survival (and a small reduction in L-P survival) at the highest caffeine concentration (1000 µg/mL) for *Df(3R)BSC852*/*Dro5^A3-B7^* animals (Fig. S2), suggesting that the *Df(3R)BSC852* deficiency fails to complement the *Dro5^A3-B7^* allele and that lesions at the Dro5 locus are indeed responsible for the increased developmental susceptibility to caffeine seen in *Dro5^A3-B7^* homozygotes (Fig. 4).

As the *Dro5^A3-B7^* allele disrupts five Dro5 EcKL genes, we aimed to test whether single-gene loss-of-function alleles for these genes would individually fail to complement the *Dro5^A3-B7^* allele for developmental survival on caffeine, which would indicate that the disrupted gene contributes to the caffeine susceptibility phenotype. Using transposable element (TE) insertion alleles for three genes—*CG31300* (Dro5-1), *CG13658* (Dro5-5) and *CG11893* (Dro5-6)—and the previously described *CG13659^38^* deletion allele as a loss-of-function allele for *CG13659* (Dro5-7), we crossed these homozygous lines to either *Dro5^A3-B7^* homozygotes or wild-type homozygotes and scored the developmental survival of their progeny on caffeine media; due to unusually high mortality on control media, we excluded these data points from fitted models (Fig. S3A,C,E). All genotypes possessing a *Dro5^A3-B7^* allele had lower L-A survival LC_50_s, suggesting a failure of the single-gene disruption alleles to complement the larger deletion (Fig. 5A, Fig. S3A); as survival of the +/*CG31300^MB00063^* and +/*CG13658^MI03110^* genotypes did not significantly respond to the change in caffeine concentration, their LC_50_s were higher than 1,000 µg/mL and therefore likely different from their corresponding *Dro5^A3-B7^* loss-of-function genotypes. P-A survival LC50s were lower for all disruption genotypes compared to their control genotypes (Fig. S3F), while L-P survival LC_50_s were only lower for *Dro5^A3-B7^*/*CG11893^MB^* and *Dro5^A3-B7^*/*CG13659^38^* animals compared to their control genotypes (Fig. S3D). These results suggest that loss of each of the four genes may contribute to the increased caffeine susceptibility of *Dro5^A3-B7^* homozygotes during the pupal stage, but only *CG11893* (Dro5-6) and *CG13659* (Dro5-7) likely contribute to the increased susceptibility during the larval stage (Table 1).

**Figure 5.**
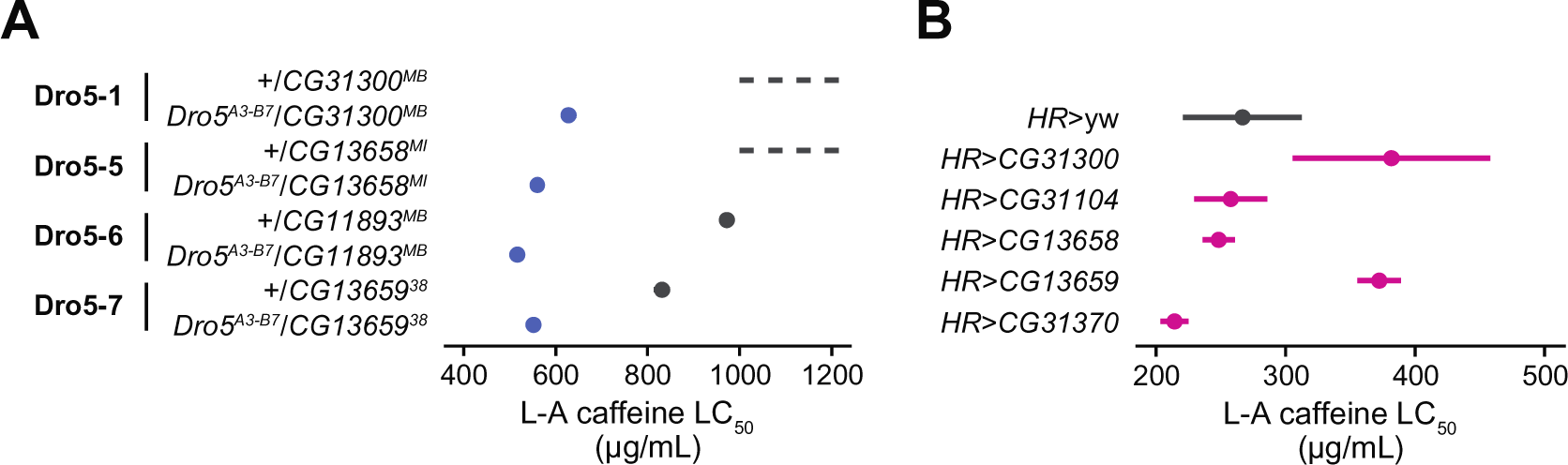
Larval–adult LC_50_ values and 95% CIs of (A) single-gene disruption animals and (B) single-gene overexpression animals on caffeine media. (A) Heterozygotes possessing a single Dro5 gene disruption allele and either a wild-type allele (+; grey) or a *Dro5^A3-B7^* allele (blue). Genotypes with a lack of dose-response effect have a dashed line to indicate an LC_50_ value above 1,000 µg/mL. For full data and LC_50_s for larval–adult, larval–pupal and pupal–adult survival, see Fig. S3. *CG31300^MB^, CG31300^MB00063^; CG13658^MI^, CG13658^MI03110^; CG11893^MB^, CG11893^MB00360^*. (B) Offspring from crossing *HR-GAL4* homozygotes and either UAS-ORF responder homozygotes for five Dro5 genes (purple) or homozygotes of the wild-type genetic background (*yw*; grey) on media containing 0–1,500 μg/mL caffeine. The *HR>CG31300* genotype was not assayed on 1,500 µg/mL media. For full data and LC_50_s for larval–adult, larval–pupal and pupal–adult survival, see Fig. S4.

### 3.5. Animals overexpressing *CG31300* (Dro5-1) and *CG13659* (Dro5-7) in detoxification tissues have increased developmental tolerance to caffeine

As a complementary test of the involvement of Dro5 EcKLs in caffeine detoxification, we misexpressed individual Dro5 UAS-ORFs using the *HR-GAL4* driver, which expresses GAL4 in the midgut, Malpighian tubules and fat body (Chung et al., 2007), and conducted dose-response developmental toxicology assays to explore if misexpression increased tolerance to caffeine compared to a control genotype (*HR*>*yw*). Unfortunately, due to stock loss, we were unable to perform these experiments with UAS-*CG11893* (Dro5-6) and UAS-*CG31436* (Dro5-10) lines. Misexpression of both *CG31300* and *CG13659* significantly increased L-A LC_50_s (Fig. 5B), with LC_50_ ratios of 1.43 (95% CI: 1.05, 1.81) and 1.40 (95% CI: 1.15, 1.65) respectively, compared to the control genotype, which was due to increased tolerance during the pupal stage (Fig. S4F) and not during the larval stage (Fig. S4D). These results are consistent with both *CG31300* and *CG13659* encoding protein products that mediate caffeine detoxification.

### 3.6. Dro5^A3-B7^ homozygotes have increased developmental susceptibility to kojic acid

In addition to caffeine, we wished to test if *Dro5^A3-B7^* mutants had increased developmental susceptibility to other naturally occurring toxins compared to wild-type animals. We chose six hydroxylated compounds: quercetin, escin, esculin, curcumin, salicin and kojic acid. For the former four compounds—soluble in ethanol—we conducted single-dose developmental toxicology assays at 40 µg/mL (quercetin) or 200 µg/mL (escin, esculin and curcumin), while for the latter two compounds— soluble in water—we conducted multiple-dose developmental toxicology assays.

*Dro5^A3-B7^* homozygotes were significantly more susceptible to kojic acid than wild-type animals, with LC_50_ ratios of 0.67 (95% CI: 0.627, 0.71), 0.79 (95% CI: 0.76, 0.82) and 0.64 (95% CI: 0.60, 0.68) for L-A, L-P and P-A survival, respectively (Fig. 6), indicating reduced tolerance at both larval and pupal stages of development. However, no significant differences in L-A, L-P or P-A survival were found between *Dro5^A3-B7^* and wild-type animals on media containing quercetin, escin, esculin or curcumin (all p > 0.05; Fig. S5). Salicin was also apparently non-toxic to both *Dro5^A3- B7^* and wild-type animals at concentrations up to 8,000 µg/mL, with no significant dose-response effect in our assays (Fig. S6). As the concentrations of quercetin, escin, esculin, curcumin and salicin used did not significantly produce development toxicity to wild-type individuals, it is possible that the doses used were not sufficient to discriminate between tolerance differences between the two genotypes, if such differences exist.

**Figure 6.**
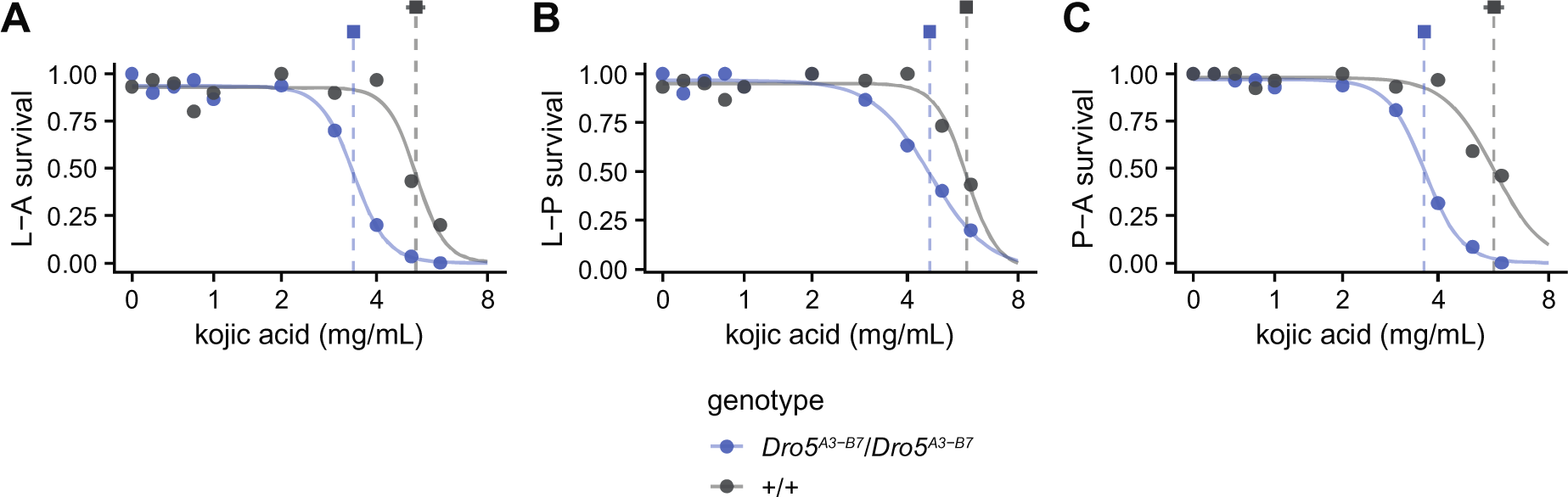
Developmental survival of +/+ homozygotes (grey) and *Dro5^A3-B7^*/*Dro5^A3-B7^* homozygotes (blue) on media containing 0–6 mg/mL kojic acid. Curves are fitted log-logistic regression dose-response models for each genotype. Dashed vertical lines and squares indicate the estimated LC_50_s for each genotype, with the horizontal bar indicating the 95% CI. (A) L-A (larval-adult) survival. (B) L-P (larval-pupal) survival. (C) P-A (pupal-adult) survival. Curves are fitted log-logistic regression dose-response models for each genotype.

We also tested if *Dro5^A3-B7^* mutants had reduced tolerance to secondary metabolites produced by *Citrus* species, decomposing fruits of which are preferred developmental substrates for *D. melanogaster* (Dweck et al., 2013). We made semi-natural fruit media with the juices of grapefruits, oranges or mandarins, and conducted developmental viability assays with homozygous *Dro5^A3-B7^* and wild-type (*Dro5^A3-B7^*/+) animals. No significant differences were found between genotypes for each medium for L-A, L-P or P-A survival (all p > 0.05, Welch’s two-sided t-test with unequal variance; Fig. S7), indicating homozygosity of the *Dro5^A3-B7^* allele does not affect developmental viability on any of the three fruit-based substrates.

## 4. Discussion

### 4.1. Genetic evidence that one or more Dro5 EcKLs in D. melanogaster confer caffeine tolerance

In this study, we have conducted the first functional experiments testing the hypothesis that members of the EcKL gene family are involved in detoxification processes in insects. Taken together, the data presented here strongly suggest that one or more EcKL genes in the Dro5 clade contribute to caffeine tolerance in *D. melanogaster* (Table 1): multiple Dro5 genes are induced by ingesting caffeine in larvae (Fig. 1D); a loss-of-function allele of *CG31370* (Dro5-8) increases adult susceptibility to caffeine in the DGRP (Fig. 2B); animals lacking five of seven Dro5 genes show decreased developmental survival on caffeine (Fig. 4); and misexpression of two Dro5 genes—*CG31300* (Dro5-1) and *CG13659* (Dro5-7)—in detoxification tissues increase developmental survival on caffeine (Fig. 6). Data showing animals lacking five Dro5 genes develop normally (Fig. 3), along with their detoxification-like transcriptional characteristics (Fig. 1; Scanlan et al., 2020), are also consistent with at least the majority of Dro5 enzymes having exogenous/xenobiotic, rather than endogenous, substrates.

*CG13659* (Dro5-7) has the strongest lines of evidence linking it to caffeine tolerance, through transcriptional induction and both knockout and misexpression toxicological phenotypes (Table 1). As *CG13659* is strongly induced by larval caffeine ingestion (Fig. 1D) and is basally expressed in the larval fat body and Malpighian tubules (Fig. 1C), this makes it likely that this gene would be involved in caffeine tolerance in wild-type animals. In contrast, while misexpression of *CG31300* (Dro5-1) reduced developmental susceptibility to caffeine, its lack of transcriptional response to caffeine (Fig. 1D), as well as its much lower basal expression in detoxification tissues (Leader et al., 2018), suggests that it is unlikely to contribute substantially to caffeine tolerance in wild-type animals. It is also possible that other Dro5 genes, such as *CG11893* (Dro5-6) and *CG31436* (Dro5-10), are involved in caffeine tolerance in wild-type animals, but due to the non-comprehensiveness of our single-gene disruption and misexpression experiments, we were unable to test this further.

It is unclear whether *CG31370* (Dro5-8) also contributes to caffeine tolerance. While the *CG31370^del^* loss-of-function allele was associated with a reduction in adult survival on caffeine media in the DGRP (Fig. 2B), misexpression of *CG31370* in detoxification tissues surprisingly decreased survival on caffeine media during larval and pupal development (Fig. 6), suggesting it does not encode an enzyme that acts in caffeine detoxification. Due to the absence of either a full Dro5-null allele or a controlled genetic background line for the *CG31370^MI07438^* TE-insertion allele, we were unable to test the developmental susceptibility of animals lacking *CG31370* function; we also did not test the tolerance of *CG31370*-misexpressing adults to caffeine. It is possible that *CG31370* encodes an enzyme that selectively acts in caffeine metabolism in adults but not pre-adult life stages; alternatively, the *CG31370^del^* allele may reduce caffeine tolerance by affecting the transcription of *CG13659*, which lies just upstream of *CG31370* (Fig. 2A) and—as previously stated—is a strong candidate for involvement in caffeine detoxification. While basal levels of expression of *CG13659* in adult females (the sex phenotyped by Najarro et al. (Najarro et al., 2015)) does not appear affected by homozygosity of *CG31370^del^*— the difference in mean log_2_(FPKM) is 0.0778 (95% CI: -0.159, 0.423) between *CG31370^wt^* and *CG31370^del^* homozygotes (Everett et al., 2020)—we hypothesise that *CG31370^del^* may affect the transcriptional induction of *CG13659* by caffeine, by disrupting a downstream transcription factor-binding site. Alternatively, *CG31370^del^* may be in linkage disequilibrium with a truly causal structural variant at the *CG13659* locus that has not yet been genotyped in the DGRP.

### 4.2. A biochemical hypothesis for EcKL-mediated caffeine detoxification by phosphorylation

The molecular targets of caffeine have been comprehensively studied in humans and other vertebrates (Fredholm et al., 1999), but the same is not true in insects— while it is known that caffeine has acute effects on the insect nervous system (Mustard, 2014), as well as chronic effects on insect development (Nathanson, 1984; Nigsch et al., 1977), the molecular causes of caffeine toxicity in *D. melanogaster* and other insects are not well understood. Molecular targets of caffeine in the insect nervous system include the ryanodine receptor and phosphodiesterases, and possibly also adenosine receptors (the main neurological target in mammals) and dopamine receptors (Mustard, 2014), some or all of which are likely responsible for caffeine’s acute effects on behaviour and physiology (Nathanson, 1984). Caffeine also inhibits proteins involved in DNA repair (Blasina et al., 1999; Tsabar et al., 2015; Zelensky et al., 2013) and increases the mutation rate *in vivo* (Kuhlmann et al., 1968), and *D. melanogaster* mutant animals with impaired genome stability are highly developmentally sensitive to caffeine (Li et al., 2013), strongly suggesting exposure to caffeine indirectly causes DNA damage *in vivo*; this mechanism is likely partially responsible for the chronic developmental toxicity of caffeine, exemplified in this study by death during metamorphosis. Feeding on food containing high concentrations of caffeine also causes death in *D. melanogaster* adults in 15–112 hours (Najarro et al., 2015), although the molecular causes of this have not been studied in detail, despite the validation of tolerance loci likely involved in detoxification (Najarro et al., 2015).

While multiple lines of evidence converge on *CG13659* conferring tolerance to caffeine in *D. melanogaster*, due to the lack of hydroxyl groups on the caffeine molecule, a kinase’s contribution to caffeine metabolism cannot be direct phosphorylation but the phosphorylation of one or more caffeine metabolites (EcKLs belong to the Group 1 kinases, which only use hydroxyl groups as a phosphoryl acceptor; Kenyon et al., 2012). As such, we propose a biochemical hypothesis for the involvement of *CG13659* and/or other EcKLs in the detoxification of caffeine (Fig. 7), which entails the existence of four classes of caffeine metabolites, all of which are produced by P450 enzymes: non-hydroxylated metabolites of much lower toxicity than caffeine; non-hydroxylated metabolites of comparable or greater toxicity than caffeine; hydroxylated metabolites of much lower toxicity than caffeine; and hydroxylated metabolites of comparable or greater toxicity than caffeine. We hypothesise that toxic hydroxylated metabolites preferentially affect DNA repair mechanisms or other targets that predominantly affect metamorphosis, and Dro5 EcKLs, such as CG13659, detoxify hydroxylated caffeine metabolites by phosphorylation, leading to a reduction in the inhibition of caffeine target proteins and increased survival on caffeine-containing media, with a bias towards conferring tolerance during metamorphosis at relatively low concentrations of caffeine (Fig. 7).

**Figure 7.**
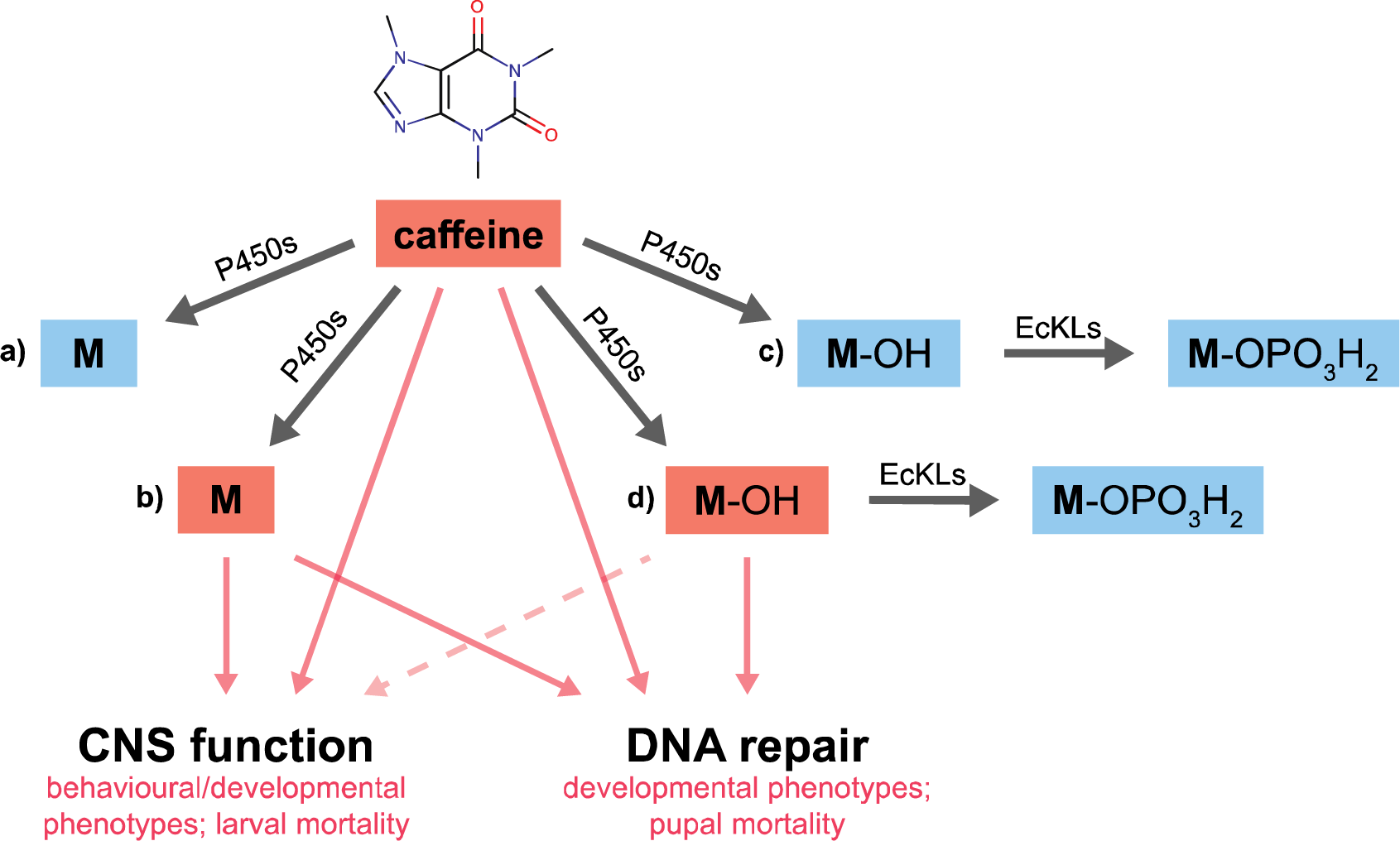
A biochemical hypothesis for the function of Dro5 EcKLs in caffeine detoxification. Ingested caffeine is metabolised by P450 enzymes to four hypothetical types of metabolites: a) non-toxic non-hydroxylated metabolites; b) toxic non-hydroxylated metabolites; c) non-toxic hydroxylated metabolites; and d) toxic hydroxylated metabolites. Hydroxylated metabolites can be phosphorylated by EcKL enzymes to form non-toxic phosphate metabolites. Toxic metabolites negatively affect (pink arrows) CNS function and/or DNA repair pathways. Toxic hydroxylated metabolites inhibit DNA repair pathways more than targets in the CNS, explaining the greater caffeine susceptibility of Dro5 mutant animals—and the greater caffeine tolerance of Dro5 overexpression animals—during the pupal stage, due to the accumulation of DNA damage in the imaginal discs, compared to the larval stages, where behavioural effects predominate. Possible complexities of caffeine metabolism wherein metabolites are acted on sequentially by multiple P450s (as suggested by Coelho et al., 2015) have not been shown, for simplicity.

The plausibility of this hypothesis is hard to judge, given that relatively little is known about caffeine metabolism in *D. melanogaster* compared to other animals. Like mammals, adult flies metabolise caffeine to the non-hydroxylated compounds theobromine, paraxanthine and theophylline through the action of P450s, but do not produce 1,3,7-trimethyluric acid, a relatively common hydroxylated metabolite of caffeine in mammals (Bonati et al., 1984). However, *D. melanogaster* also produces an additional five unidentified metabolites, one of which—M2—is the second-most abundant caffeine metabolite in male flies two hours after exposure, accounting for 34% of ingested caffeine (Coelho et al., 2015), suggesting its formation may be important for reducing the toxicity of caffeine; it is possible one or more of these unidentified metabolites are hydroxylated. Based on RNAi knockdown and radiolabelling experiments, Coelho *et al*. (2015) hypothesised that the M2 metabolite is produced by Cyp12d1 and then subsequently metabolised by one or both of Cyp6a8 and Cyp6d5, while theobromine is produced by Cyp6d5 and metabolised by Cyp6a8; independently, QTL mapping and RNAi experiments by Najarro *et al*. (2015) indicated that both *Cyp12d1* and *Cyp6d5* contribute to caffeine tolerance in adult flies. Taken together, these data suggest that the formation and/or further metabolism of M2 and theobromine, which together account for ∼76% of metabolised caffeine (Coelho et al., 2015), strongly influence the tolerance of *D. melanogaster* to caffeine exposure. The presence of hydroxyl groups on any significantly abundant caffeine metabolites in *D. melanogaster* would produce a plausible substrate for Dro5 enzymes, including CG13659.

Additionally, we recently found that a non-synonymous variant (W260S) in another P450, *Cyp4s3*, is associated with developmental caffeine survival in the DGRP (Scanlan et al., 2020), based on a reanalysis of phenotype data from Montgomery *et al*. (2014); despite *Cyp4s3* not being induced by caffeine exposure in larvae or adults (Coelho et al., 2015; Willoughby et al., 2006), it may also be involved in caffeine metabolism, although its possible role is unclear.

Our biochemical hypothesis for the action of Dro5 EcKLs in caffeine detoxification relies on the sustained toxicity of hypothetical hydroxylated caffeine metabolites. While caffeine metabolites and other methylxanthines can have physiological effects in humans sometimes equal or exceeding that of caffeine itself (Benowitz et al., 1995; Geraets et al., 2006; Malki et al., 2006), only limited data currently exist on the differential toxicity of caffeine and its metabolites in insects: theobromine appears less toxic than caffeine in *D. melanogaster* adults (Matsagas et al., 2009); while caffeine, theophylline and theobromine are toxic to the pupal CNS of giant silkmoths (Lepidoptera: Bombyoidea); caffeine and theophylline are 3- to 4-fold more toxic than theobromine (Blaustein and Schneiderman, 1960); and theophylline and theobromine are not toxic at daily doses of 5–10 µg in *Vespa orientalis* (Hymenoptera: Vespoidea) and *Apis mellifera* (Hymenoptera: Apoidea), unlike caffeine (Ishay and Paniry, 1979). Additionally, despite being a canonical phase I detoxification reaction, hydroxylation can bioactivate some pro-toxic xenobiotic compounds (Harrop et al., 2018; Idda et al., 2020; Salgado and David, 2017). As the metabolism of caffeine is poorly understood at a fine level of detail even in model insect species like *D. melanogaster*, the relative change in toxicity of caffeine at each step of its metabolism in insects remains to be determined.

### 4.3. How ecologically relevant is caffeine detoxification in D. melanogaster?

Caffeine is found in the leaves, fruits, seeds and/or flowers of a variety of plants, including species in the genera *Coffea*, *Camellia*, *Theobroma*, *Paullinia*, *Cola*, *Ilex* and *Citrus* (Anaya et al., 2006) and is primarily thought to be an antifeedant against invertebrate herbivores (Hollingsworth et al., 2002; Nathanson, 1984; Uefuji et al., 2005), although it can also function to enhance pollinator learning and memory, improving foraging rates (reviewed by Stevenson, 2020). *D. melanogaster* is a saprophage that feeds on rotting fruit substrates (Markow, 2019), which are unlikely to originate from the small, caffeine-rich fruits found in the *Coffea*, *Cola* and *Paullinia* genera. However, *Citrus* fruits produce highly favourable substrates for *D. melanogaster* (Dweck et al., 2013)—while caffeine is found in *Citrus* flowers, not fruits (Kretschmar and Baumann, 1999), *Citrus* trees typically produce large numbers of flowers (Iglesias et al., 2007), raising the possibility that fruits and flowers decompose together, forming a food substrate for *D. melanogaster* containing toxicologically relevant levels of caffeine. Whole *Citrus* flowers contain approximately 318 nmol/g (62 μg/g) caffeine (Kretschmar and Baumann, 1999), meaning that a 1:1 flower to fruit ratio—a plausible upper limit for what might be found in nature—would produce a developmental substrate with 31 μg/g caffeine. This is below the developmental LC_50_s determined for wild-type animals in this study but might produce adverse behavioural or developmental effects in natural environments, especially for non-adapted genotypes, producing selection for efficient caffeine detoxification. A diverse collection of *Drosophila* species other than *D. melanogaster* use *Citrus* spp. fruits as developmental substrates in nature (Hoenigsberg et al., 1977), suggesting the ability to detoxify caffeine and related methylxanthine compounds may have been present in the ancestor of all or most Drosophila species.

Alternatively, the ability to detoxify caffeine may be due to generalist detoxification mechanisms, possibly related to those metabolising other alkaloids unlikely to be in the natural diet of *D. melanogaster*, such as nicotine, carnegine and isoquinoline alkaloids (Danielson et al., 1995; Fogleman, 2000; Highfill et al., 2017; Marriage et al., 2014).

### 4.4. EcKL-mediated tolerance of kojic acid and other toxins

The dramatic and substantial expansion of the Dro5 clade in the *Drosophila* genus (Scanlan et al., 2020) is suggestive of a role in detoxification processes relevant to the ecological niches of this group of largely saprophagous insects (Markow, 2019). In this study, we found preliminary evidence that Dro5 EcKLs confer tolerance to the hydroxylated fungal secondary metabolite kojic acid (Fig. 6)—however, we did not perform further experiments to dissect which gene or genes disrupted in *Dro5^A3-B7^* homozygotes may be responsible. We decided to use kojic acid in our experiments because it is both toxic to *D. melanogaster* (Dobias et al., 1977) and produced as a secondary metabolite of known filamentous fungal competitors of *Drosophila* larvae (El-Kady et al., 2014; Rohlfs et al., 2005). While the concentrations of kojic acid used were relatively high (up to 6 mg/mL or 0.6% w/v), they are likely to be ecologically relevant, as many strains of *Aspergillus* spp. and *Penicillium* spp. regularly produce more than 0.5% w/v kojic acid in culture (Beard and Walton, 1969; El-Kady et al., 2014). Given this, it is likely that *D. melanogaster* and other *Drosophila* spp. have evolved metabolic detoxification mechanisms to increase their tolerance to kojic acid. Essentially nothing is known about the metabolism of kojic acid in insects, although it is substantially metabolised to sulfate and glucuronide conjugates in rats (Burnett et al., 2010), suggesting similarly conjugation-heavy metabolism—such as phosphorylation via EcKLs—could also occur in insects.

Other fungal secondary metabolites that are plausible substrates for EcKLs are the hydroxylated mycotoxins citrinin and patulin, which are synthesised by various *Aspergillus* and *Penicillium* species (Paterson et al., 1987; Puel et al., 2010). Of note, *CG31104* (Dro5-2), *CG11893* (Dro5-6) and *CG13659* (Dro5-7) are all transcriptionally induced by feeding on wild-type vs. secondary metabolite-deficient strains of *Aspergillus nidulans* (Trienens et al., 2017), raising the possibility that one or more defensive compounds produced by *A. nidulans* specifically could also be substrates for Dro5 EcKLs.

We did not find evidence that Dro5 genes contribute to tolerance to the plant secondary metabolites quercetin, esculin, escin, curcurmin and salicin, possibly due to the use of indiscriminate toxin concentrations (Figs. S6 & S7). Salicin was an attractive compound for use in this study because it is phosphorylated by *Lymantria dispar* (Lepidoptera: Noctuoidea), along with four similar glycosides—arbutin, helicin, phenol glycoside and catechol glucoside (Boeckler et al., 2016). Cyanogenic glucosides present in cassava (*Manihot esculenta*) were also recently found to be phosphorylated by the silverleaf whitefly, *Bemisia tabaci* (Hemiptera: Aleyrodoidea; Easson et al., 2021), glycosidic metabolites of the drug midazolam are phosphorylated in the locust *Schistocerca gregaria* (Orthoptera: Acridoidea; Olsen et al., 2015), and phosphorylated glycosides are also formed by other species in the orders Blattodea, Coleoptera, Dermaptera, Diptera and Lepidoptera (Ngah and Smith, 1983). The phosphorylation of glycosides has been hypothesised to inhibit hydrolysis post-ingestion, preventing the formation of toxic aglycones (Boeckler et al., 2016)—indeed, phosphorylated linamarin metabolites cannot be hydrolysed to cyanogenic aglycones by *B. tabaci* transglucosidases *in vitro* (Easson et al., 2021), suggesting that phosphorylation can act directly on toxins before other metabolic reactions have occurred. It is currently unknown if *D. melanogaster* can phosphorylate xenobiotic glycosides, although glycosides—particularly those of flavonoids—are abundant secondary metabolites present in the fruits of *Citrus* spp. (Wang et al., 2017). However, in this study we did not find evidence for differences in the tolerance of wild-type and *Dro5^A3-B7^* mutant animals feeding on semi-natural *Citrus* fruit media (Fig. S7), suggesting either disruption of the five EcKL genes is not sufficient to significantly affect tolerance to the mixture of compounds present in these diets, or that these genes do not contribute to tolerance of *Citrus* spp. secondary metabolites at all.

Another possible substrate for Dro5 EcKLs, or indeed any detoxification-candidate EcKLs in *D. melanogaster*, is harmol, the only xenobiotic compound known to be phosphorylated in this species (Baars et al., 1980). Harmol is a human metabolite of harmine, a harmala alkaloid found in the ayahuasca plant *Banisteriopsis caapi* (Riba et al., 2003), and may not be found in the natural diet of *D. melanogaster*, although harmine appears only mildly developmentally toxic up to concentrations of at least 200 µg/mL (Cui et al., 2020), suggesting it is efficiently detoxified, possibly through a metabolic pathway that includes phosphorylation.

We note that the *D. melanogaster* EcKL gene *CHKov1* (Dro18-1) has been reported as associated with resistance to the organophosphate (OP) insecticide azinphos-methyl, supported by backcrossing a TE-insertion allele into a wild-type genetic background and conducting adult survival assays (Aminetzach et al., 2005), but a larger study using a developmental survival phenotype in the DGRP failed to find a significant association between the *CHKov1* locus and OP resistance (Battlay et al., 2016); however, TE-insertion and further-derived duplication alleles at *CHKov1* have been convincingly linked to viral resistance (Magwire et al., 2012, 2011). Regardless of the phenotypes associated with this locus, the alleles in question substantially disrupt the coding region of *CHKov1* and are therefore unlikely to reflect the native functions of EcKL enzymes.

### 4.5. Alternative hypotheses for results, study limitations and future research questions

An alternative hypothesis explaining the results in this paper is that Dro5 genes are involved in a general tolerance process to toxins and are not directly involved in detoxification *per se*. This was hypothesised for the paralogous EcKLs *CG16898* (Dro26-1) and *CG33301* (Dro26-2), variants near which are associated with resistance to multiple, chemically unrelated toxic stresses (Scanlan et al., 2020). If this alternative hypothesis is true, it may point towards the existence of an undiscovered generalist toxin-response pathway in *D. melanogaster*, as Dro5 genes (and Dro26-1 and Dro26-2) are not transcriptionally regulated by the ROS-sensitive CncC pathway (Misra et al., 2011), yet respond to the ingestion of many different xenobiotic compounds (Scanlan et al., 2020). Curiously, *CG13659*, *CG11893* and *CG16898* (but not other EcKLs) may be positively regulated in larvae by XBP1 (Huang et al., 2017), an evolutionarily conserved physiological stress-response transcription factor that mediates sensitivity to oxidative stress (Liu et al., 2009) and may be a good candidate for mediating toxin responses in *D. melanogaster*.

This study is limited by the non-comprehensive nature of some of our experiments, such as a lack of data on the changes in caffeine susceptibility upon *CG11893* (Dro5-6) and *CG31436* (Dro5-10) misexpression, as well as a lack of single-gene disruption data for *CG31104* (Dro5-2), *CG31370* (Dro5-8) and *CG31436*, as well as the lack of a full seven-gene Dro5 null allele. Some toxicological assays were also conducted with a limited caffeine concentration range, which should be replicated in the future to properly calculate LC_50_s. This study is also limited by its use of genetic experiments alone to test a detoxification hypothesis, which ideally should be done through a combination of genetic, toxicological and biochemical experiments. Radiolabelled or isotope-labelled caffeine metabolite tracing, combined with Dro5 gene knockout or misexpression, should determine if phosphate conjugates of caffeine metabolites are indeed produced by Dro5 enzymes, an approach that could be complemented with *in vitro* studies of Dro5 enzyme activity and/or structure.

Future work could also focus on adult caffeine susceptibility and whether it is altered by any of the genetic manipulations described in this study; the caffeine susceptibility of the homozygous *CG31370^del^* genotype also needs to be validated through further toxicological experiments. More tissue-specific misexpression and knockout experiments, as well as explorations of tissue-specific induction by caffeine exposure, could improve our understanding of how and where *CG13659* (Dro5-7) confers caffeine tolerance. Further experiments are also clearly needed to explore the relationship between Dro5 EcKLs and kojic acid, as well as other fungal secondary metabolites and ecologically relevant toxins for *D. melanogaster*, which could be probed with fungal-larval competition assays ala. Trienens *et al*. (2010).

## 5. Summary

This study has provided the first experimental evidence that insect EcKL genes are involved in detoxification in the model insect *Drosophila melanogaster*. Multiple lines of evidence have linked the Dro5 genes—a large, dynamic clade containing many detoxification candidate genes (Scanlan et al., 2020)—to tolerance of the plant alkaloid caffeine, and suggest an additional association with the fungal secondary metabolite kojic acid, both of which may be ecologically relevant toxins for *D. melanogaster*. This work lays the groundwork for future research into detoxicative kinases and may lead to a deeper understanding of caffeine metabolism in insects.

## Abbreviations

ABC: ATP-binding cassette
CNS: Central nervous system
DGRP: Drosophila Genetic Reference Panel
GWAS: Genome-wide association study
EcKL: Ecdysteroid kinase-like
UDP: Uridine diphosphate

## Author contributions

**Jack L Scanlan**: Conceptualization, Methodology, Formal Analysis, Investigation, Writing – Original Draft, Writing – Review & Editing, Visualization.

**Paul Battlay**: Methodology, Formal Analysis, Writing – Review & Editing.

**Charles Robin**: Conceptualization, Writing – Review & Editing, Supervision, Project administration.

## Acknowledgments

The authors would like to thank Simon Bullock for the pCFD6 plasmid, the Drosophila Genomics Resource Center for the UAS-ORF plasmids, Jin Kee for initial insights into transcriptomic data, and Philip Batterham and Trent Perry for *Drosophila* stocks.

## Supplementary Materials

### Supplementary Figures

**Figure S1.**
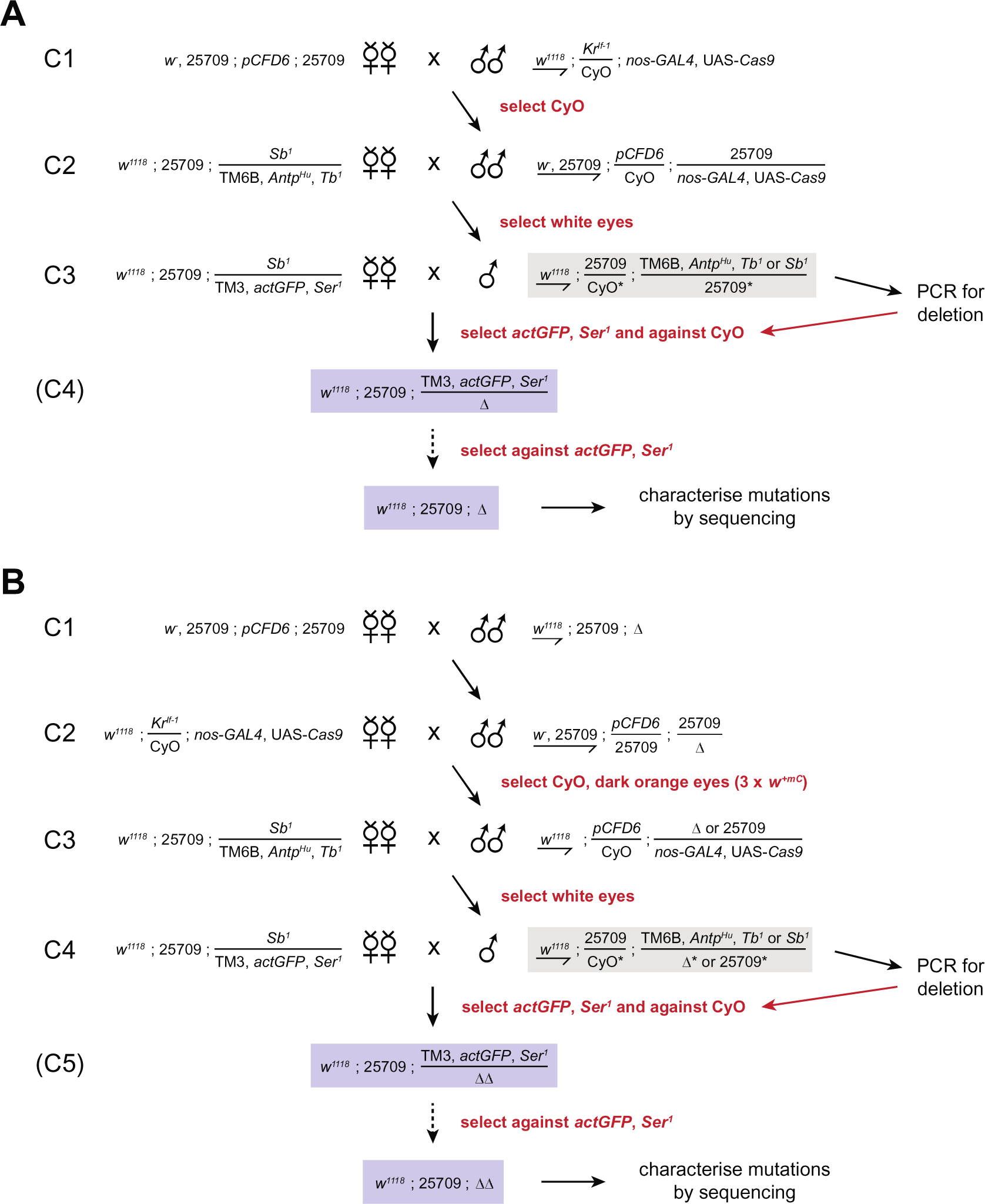
Crossing schemes for CRISPR-Cas9 mutagenesis on chromosome 3 (chr3) using *pCFD6*-transformed flies. (A) Initial mutagenesis of a wild-type chr3 locus in the BL25709 genetic background to produce a deletion allele (Δ). The single males used in C3 (grey box) are ‘founder males’ of each potential mutant line. (B) Mutagenesis of an already-mutagenised chr3 (Δ) to produce a double mutant line (ΔΔ). In this scheme, founder males (grey box) are the single males used in C4. Males used in C3 are selected by the colour of their eyes—as the *pCFD6*, *nos-GAL4* and UAS-*Cas9* constructs all contain a mini-white gene (*w^+mC^*) that produces orange eyes in a *w^-^* background, individuals that inherit all three transgenic constructs (i.e. mutagenic males) can be distinguished from those that only inherit only two (*nos-GAL4* and UAS-*Cas9*). The PCR step after C4 (scheme B) needs to check for the presence of both the initial mutation as well as any new mutations produced. Dashed arrows indicate a possible homozygosing step (if the alleles generated are homozygous-viable). C3 in (A) and C4 in (B) can use either TM3, *actGFP*, *Ser^1^* or *Sb^1^* males as founders, in order to double the number of potential mutant lines that can be generated from the cross.

**Figure S2.**
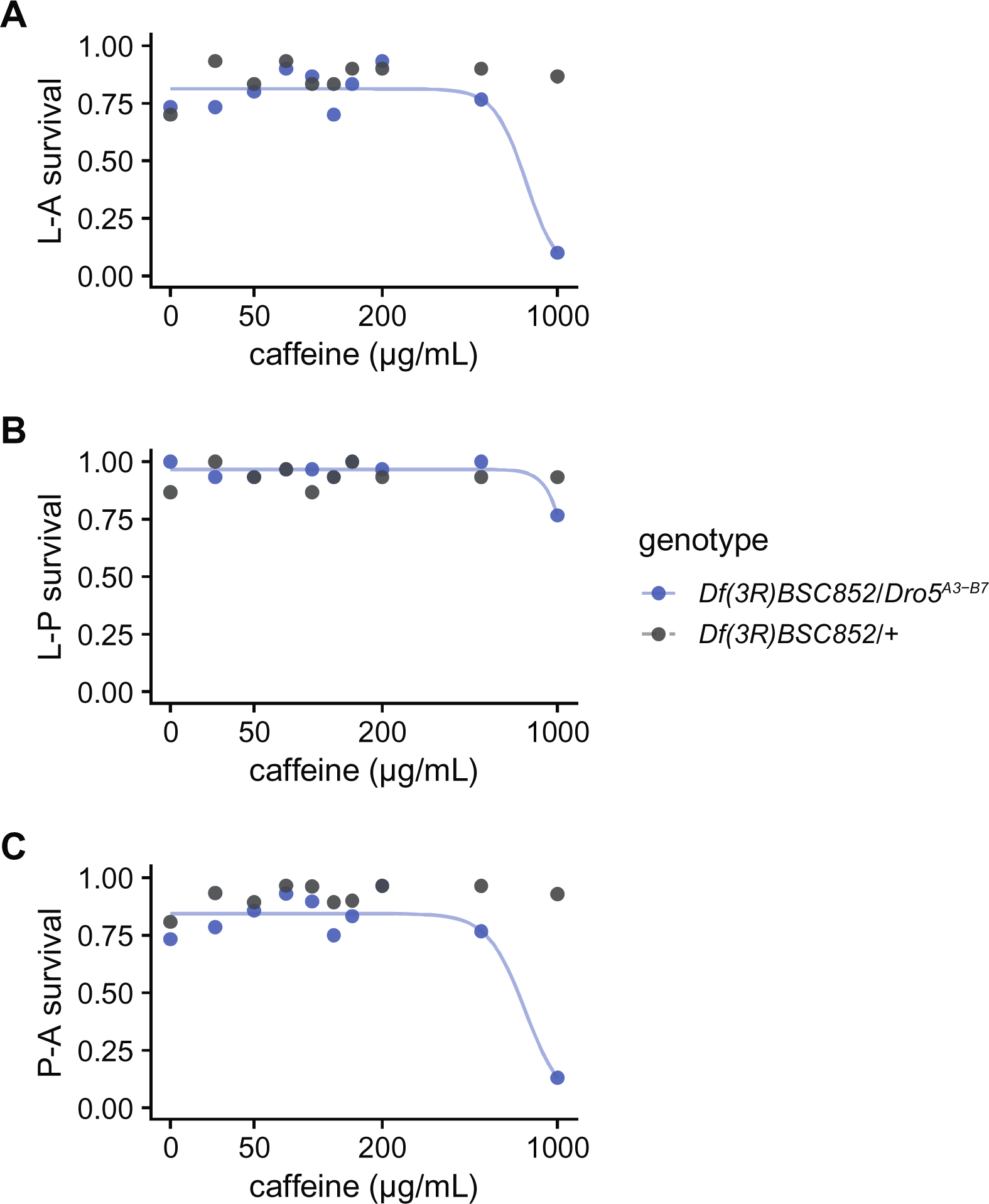
Developmental survival of *Df(3R)BSC852*/*Dro5^A3-B7^* animals (blue) and *Df(3R)BSC852*/+ animals (grey) on media containing 0-1,000 μg/mL caffeine. Curves are fitted log-logistic regression dose-response models for each genotype, excluding survival on the control media (0 µg/mL, open circles). Where data did not significantly contain a dose-response effect, no curve has been fitted. (A) Larval-adult (L-A) survival. (B) Larval-pupal (L-P) survival. (C) Pupal-adult (P-A) survival.

**Figure S3.**
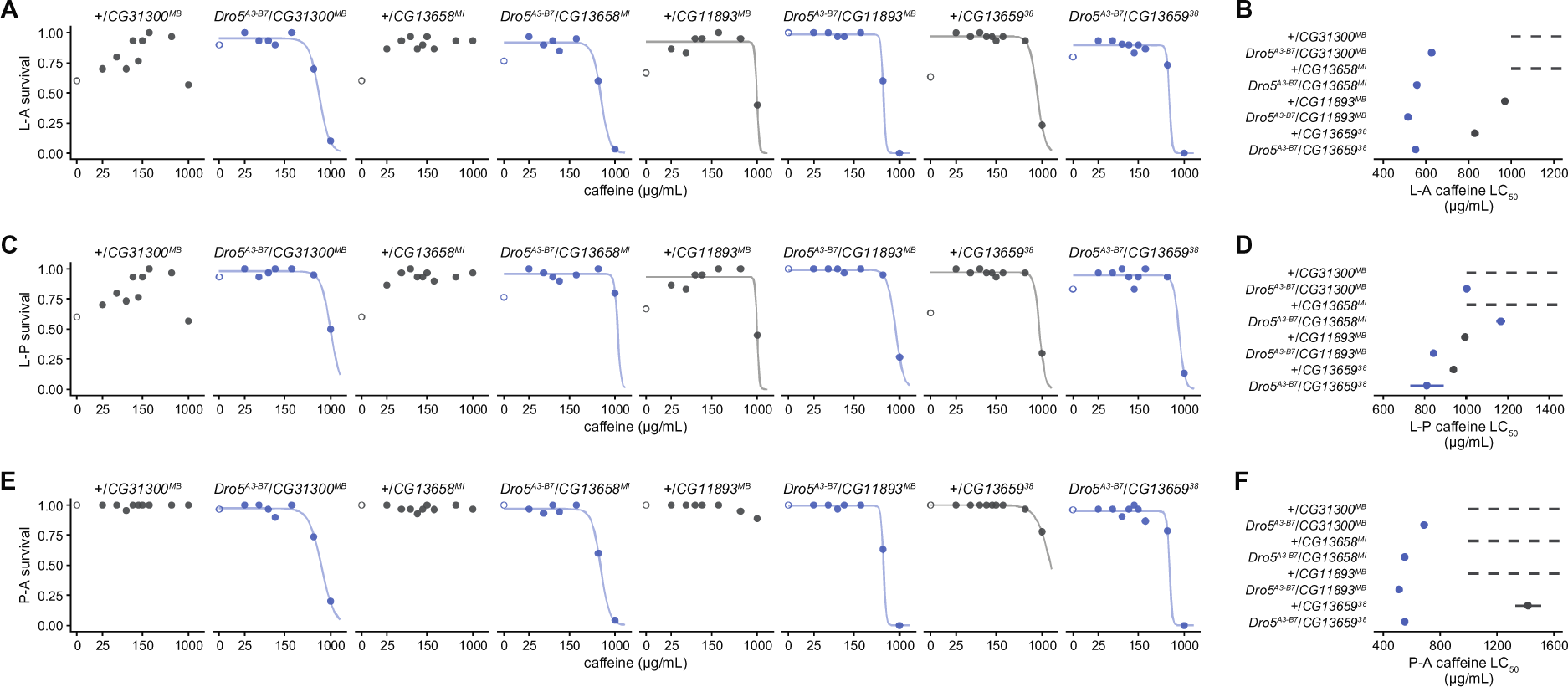
Developmental survival of heterozygotes of single-gene disruption alleles and the wild-type allele (+, grey) and heterozygotes of single-gene disruption alleles and the *Dro5^A3-B7^* allele (blue) on media containing 0-1,000μg/mL caffeine. L-A, larval-adult; L-P, larval-pupal; P-A, pupal-adult. (A,C,E) Proportional survival for each of the three developmental survival types. Curves are fitted log-logistic regression dose-response models for each genotype, excluding survival on the control media (0 µg/mL, open circles). Where data did not significantly contain a dose-response effect, no curve has been fitted. (B,D,F) LC_50_ values and 95% CIs on caffeine media for each of the three survival types, for each genotype. Genotypes without fitted models (due to a lack of dose-response effect) have a dashed line to indicate a likely LC_50_ value above 1,000 µg/mL.

**Figure S4.**
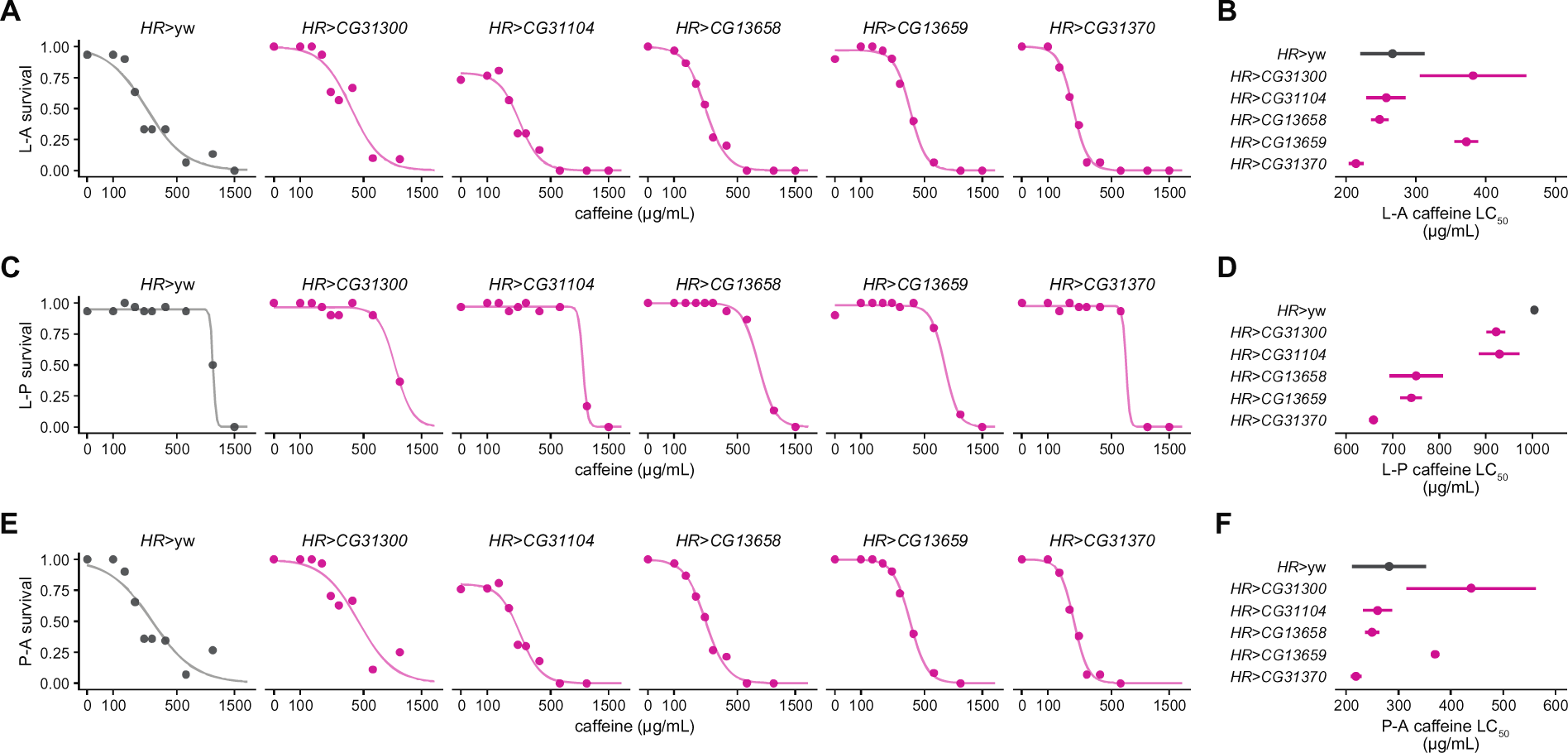
Developmental survival of offspring from crossing *HR-GAL4* homozygotes and either UAS-ORF responder homozygotes for five Dro5 genes (purple) or homozygotes of the wild-type genetic background (*yw*, grey) on media containing 0–1,500 μg/mL caffeine. The *HR>CG31300* genotype was not assayed on 1,500 µg/mL media. L-A, larval-adult; L-P, larval-pupal; P-A, pupal-adult. (A,C,E) Proportional survival for each of the three developmental survival types. Curves are fitted log-logistic regression dose-response models for each genotype. (B,D,F) LC_50_ values and 95% CIs on caffeine media for each of the three survival types, for each genotype.

**Figure S5.**
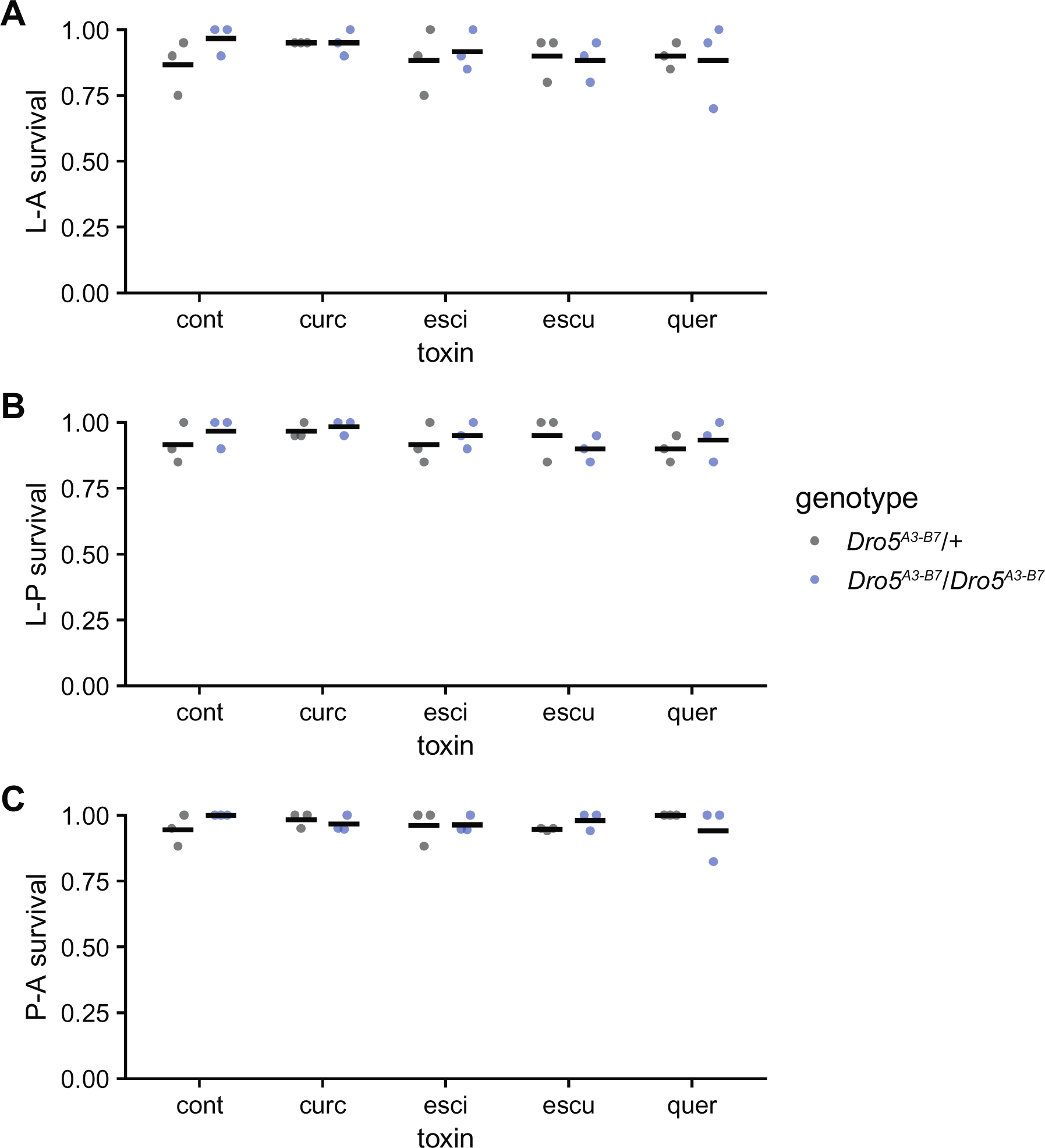
Developmental survival of *Dro5^A3-B7^*/*Dro5^A3-B7^* animals (blue) and *Dro5^A3-B7^*/+ animals (grey) on media containing curcumin (curc; 200 µg/mL), escin (esci; 200 µg/mL), esculin (escu; 200 µg/mL) or quercetin (40 µg/ml), or control media (cont; EtOH only). Each dot is a vial of 20 larvae; black bars indicate the mean. (A) Larval-adult (L-A) survival. (B) Larval-pupal (L-P) survival. (C) Pupal-adult (P-A) survival.

**Figure S6.**
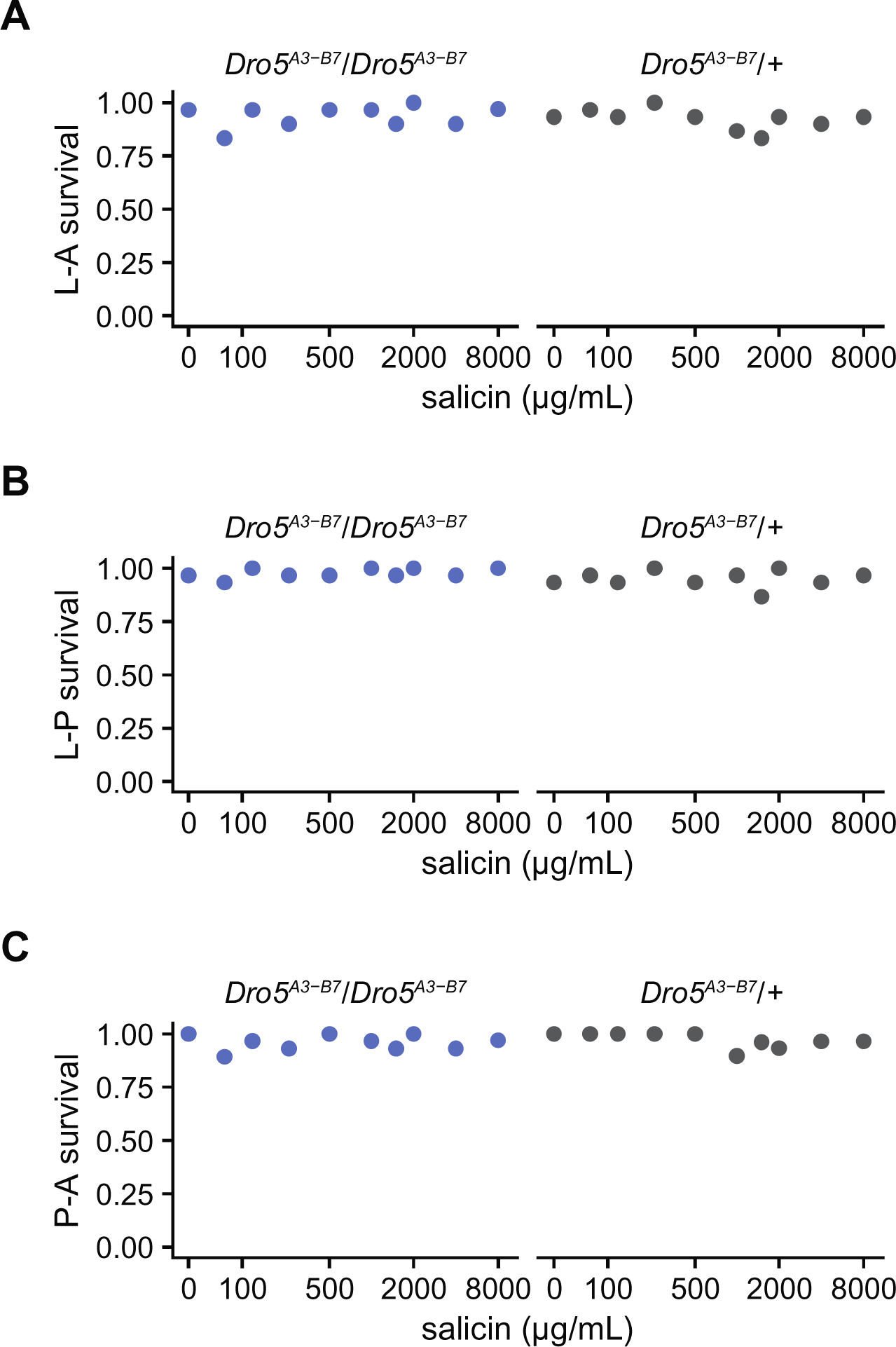
Developmental survival of *Dro5^A3-B7^*/*Dro5^A3-B7^* animals (blue) and *Dro5^A3-B7^*/+ animals (grey) on media containing 0-8,000 μg/mL salicin. No dose-response models have been fitted due to a lack of significant dose-response effects. (A) Larval-adult (L-A) survival. (B) Larval-pupal (L-P) survival. (C) Pupal-adult (P-A) survival.

**Figure S7.**
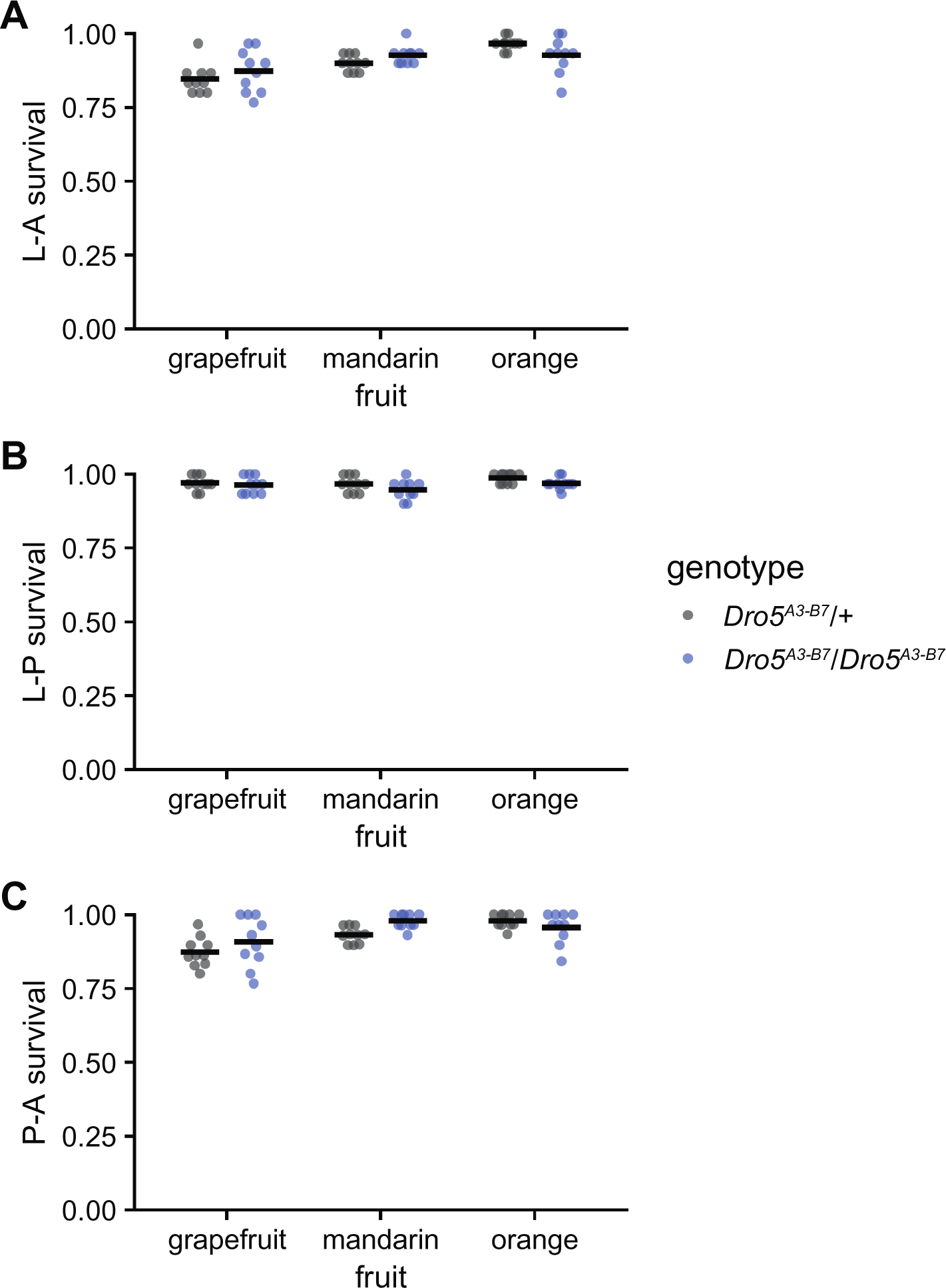
Developmental survival of *Dro5^A3-B7^*/*Dro5^A3-B7^* animals (blue) and *Dro5^A3-B7^*/+ animals (grey) on media containing grapefruit, mandarin or orange juice. Each dot is a vial of 30 larvae; black bars indicate the mean. (A) Larval-adult (L-A) survival. (B) Larval-pupal (L-P) survival. (C) Pupal-adult (P-A) survival.

### Supplementary Tables

**Table S1.**
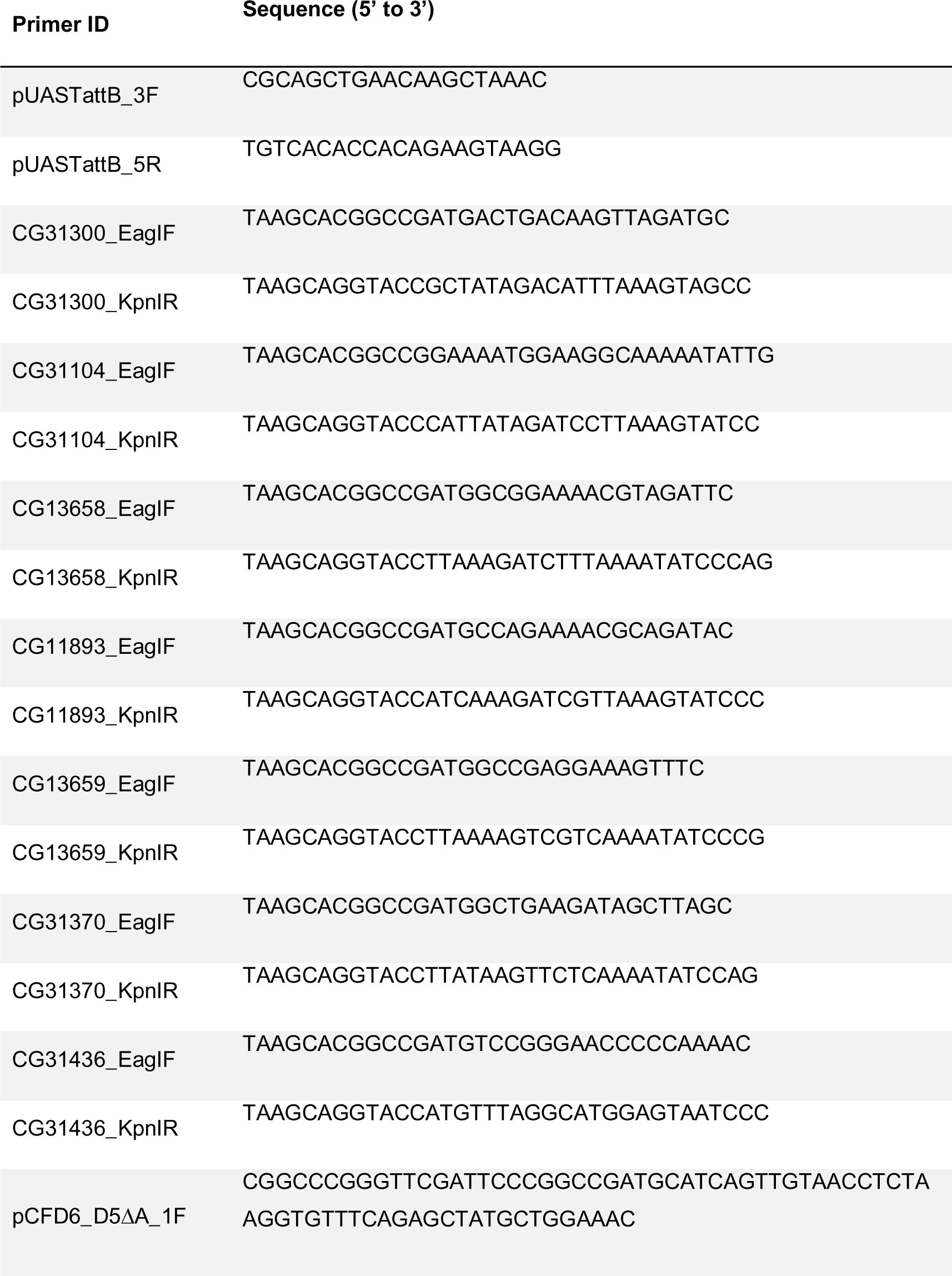
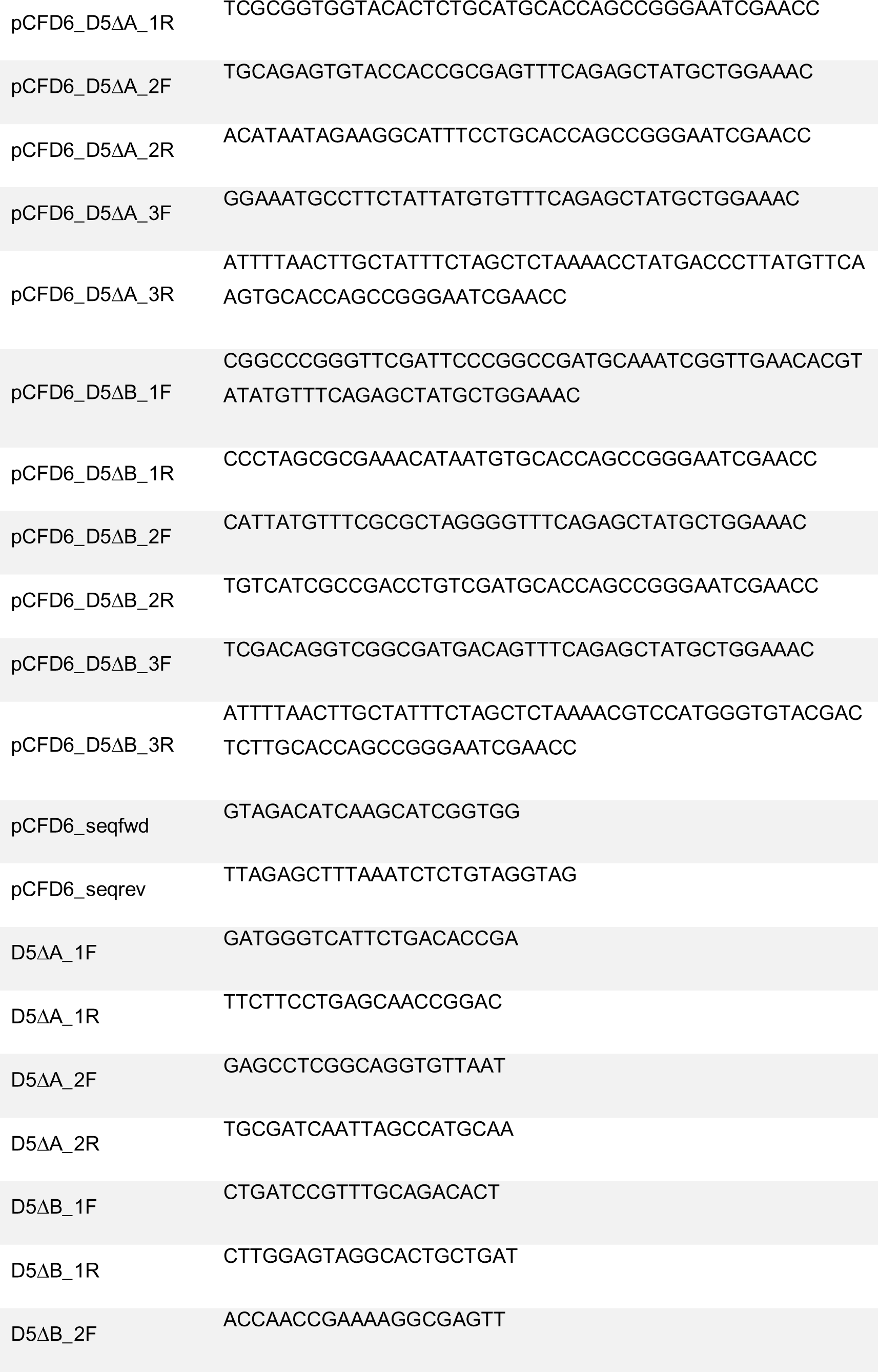
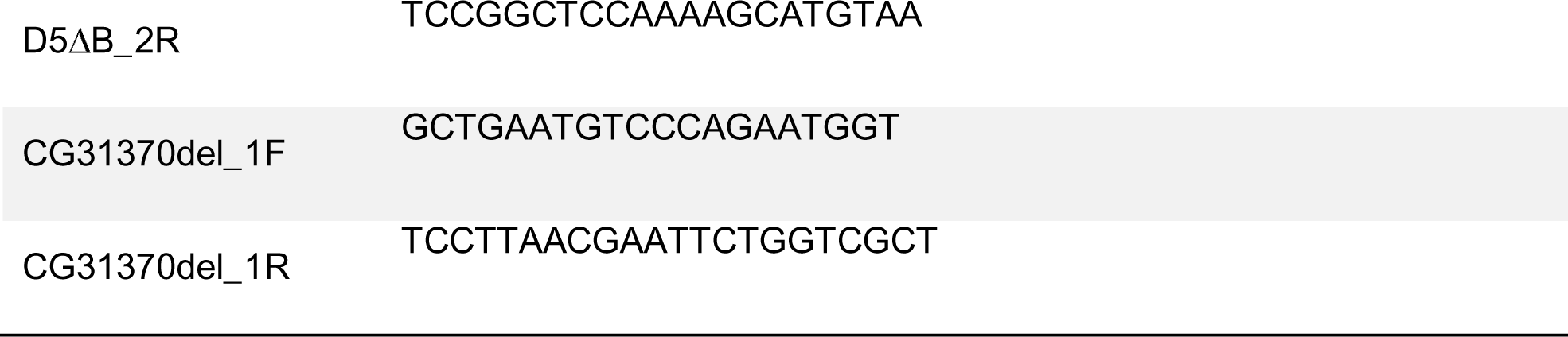
List of primer sequences used for cloning and genotyping.

**Table S2.** DGRP genotypes for *CG31370* and caffeine tolerance phenotypes. [See file “Table S2.xlsx”]

