## Supplementary figures and images for "Ecdysteroid kinase-like (EcKL) paralogs confer developmental tolerance to caffeine in *Drosophila melanogaster*"

### Figure 1

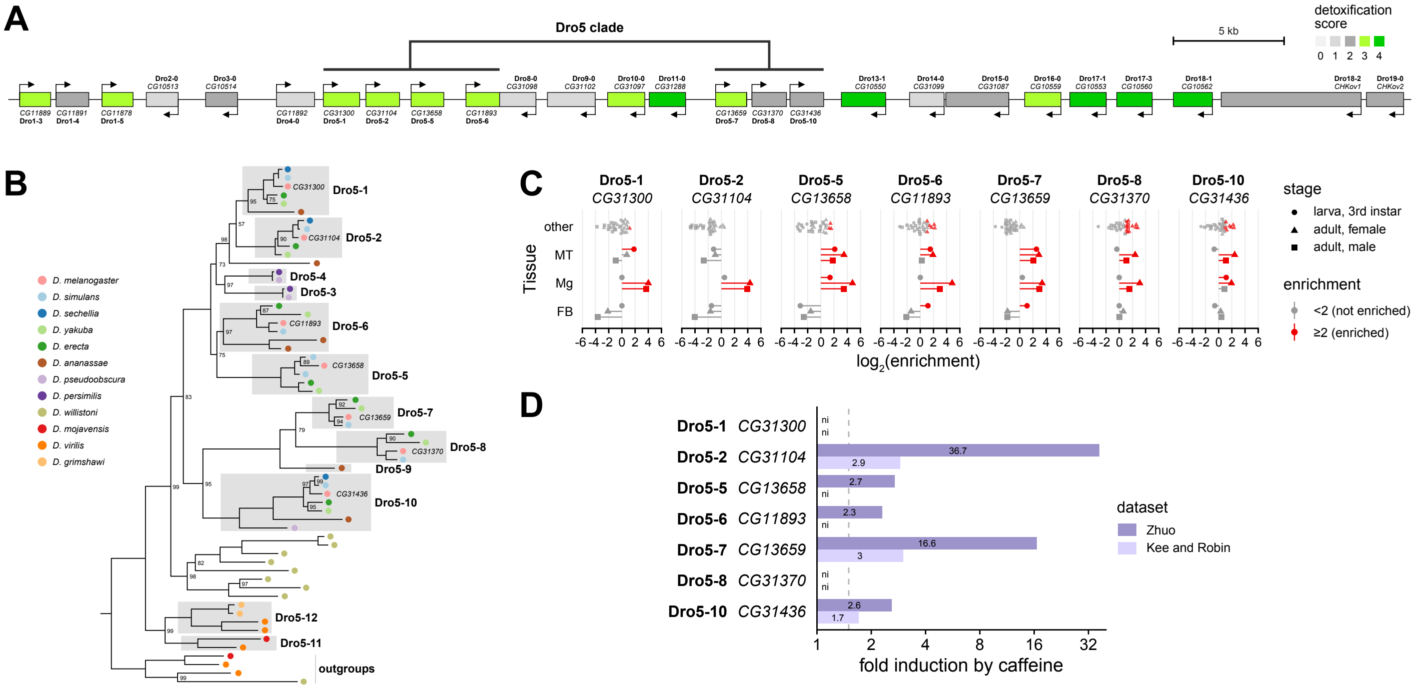

### Figure 2

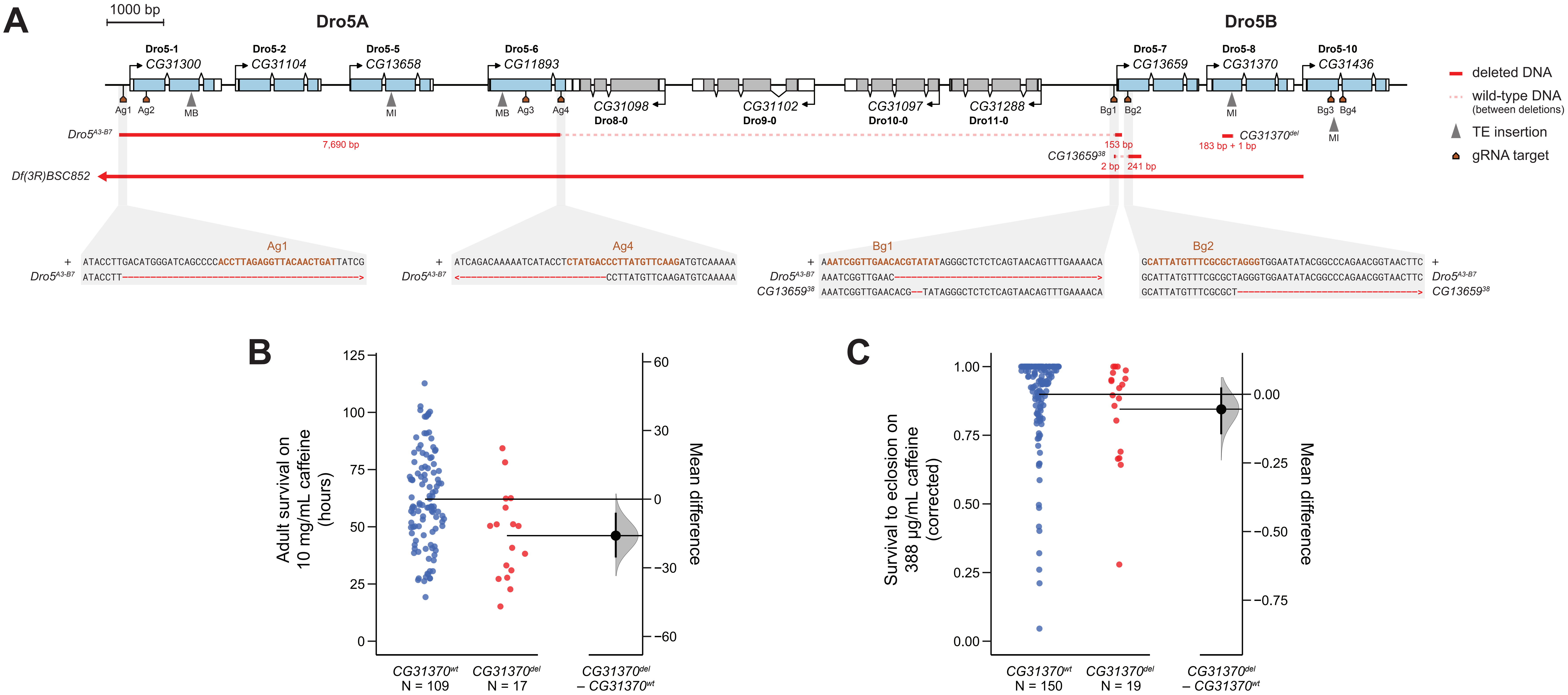

### Figure 3

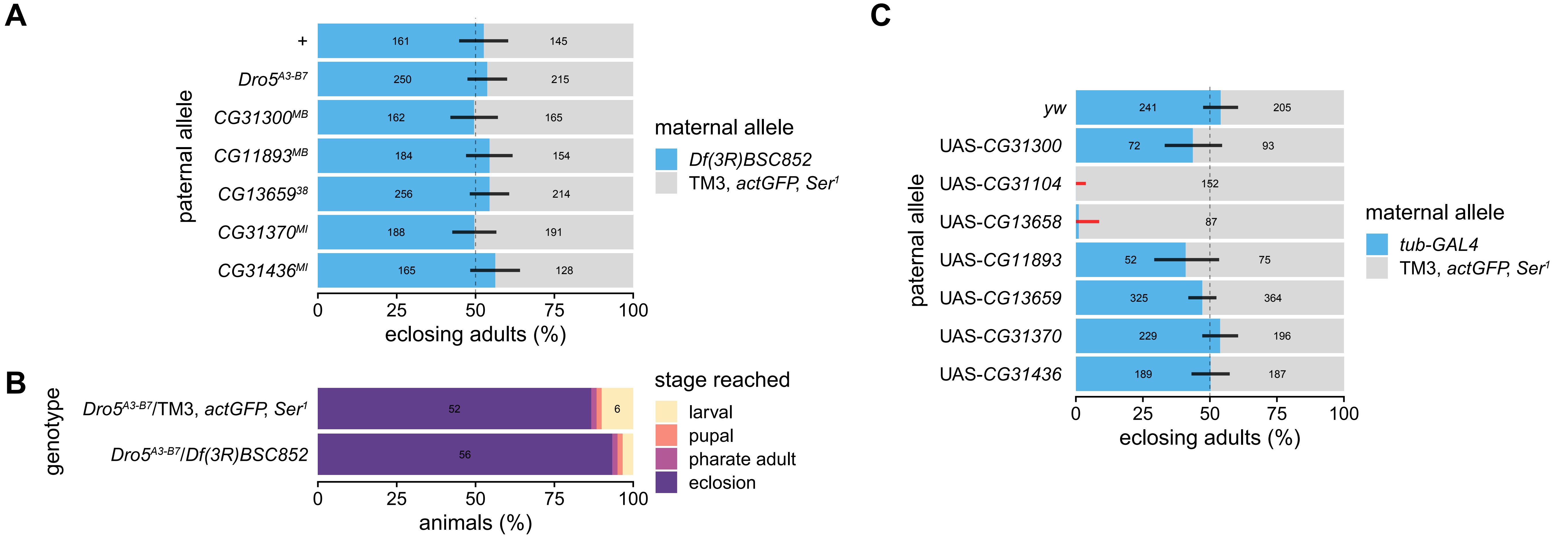

### Figure 4

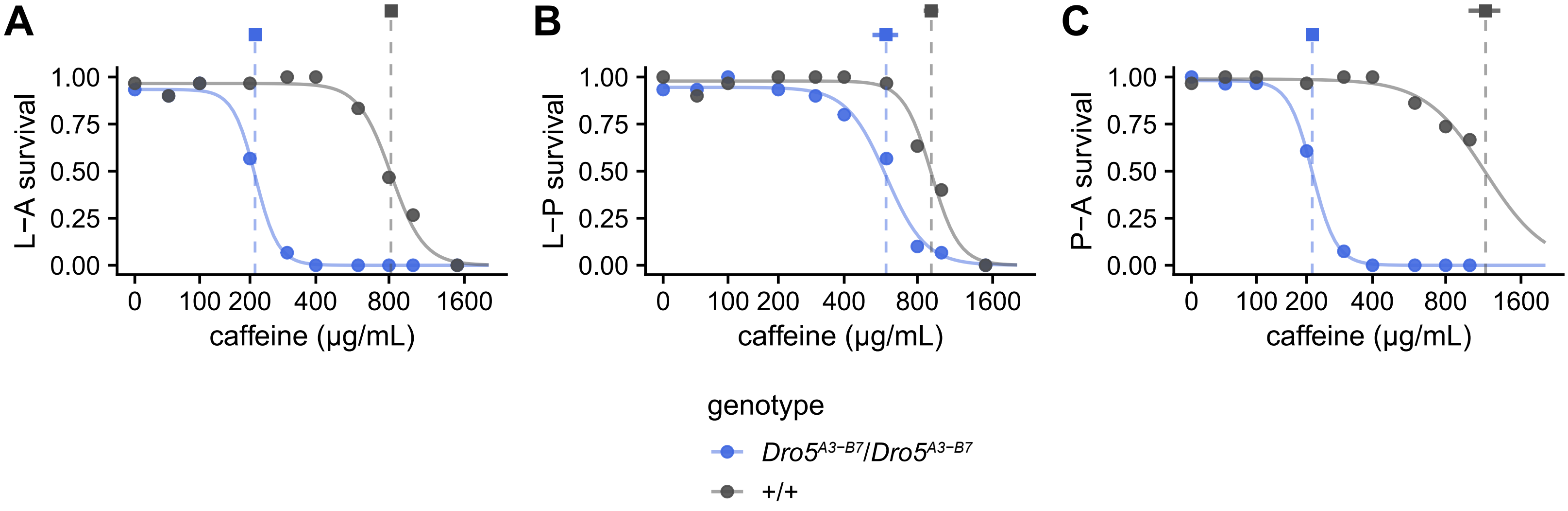

### Figure 5

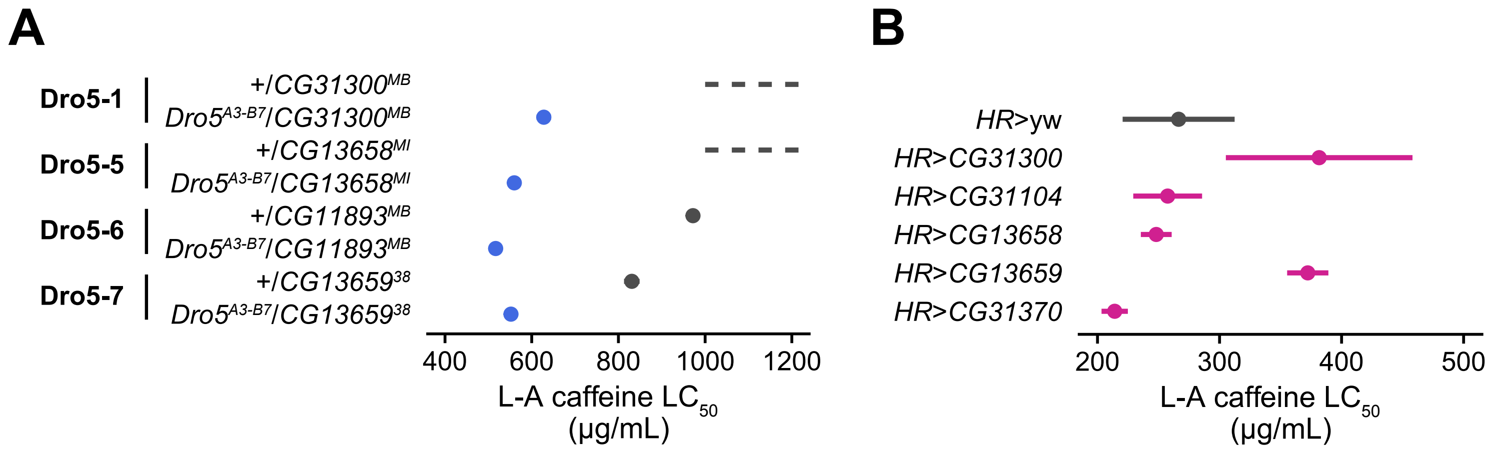

### Figure 6

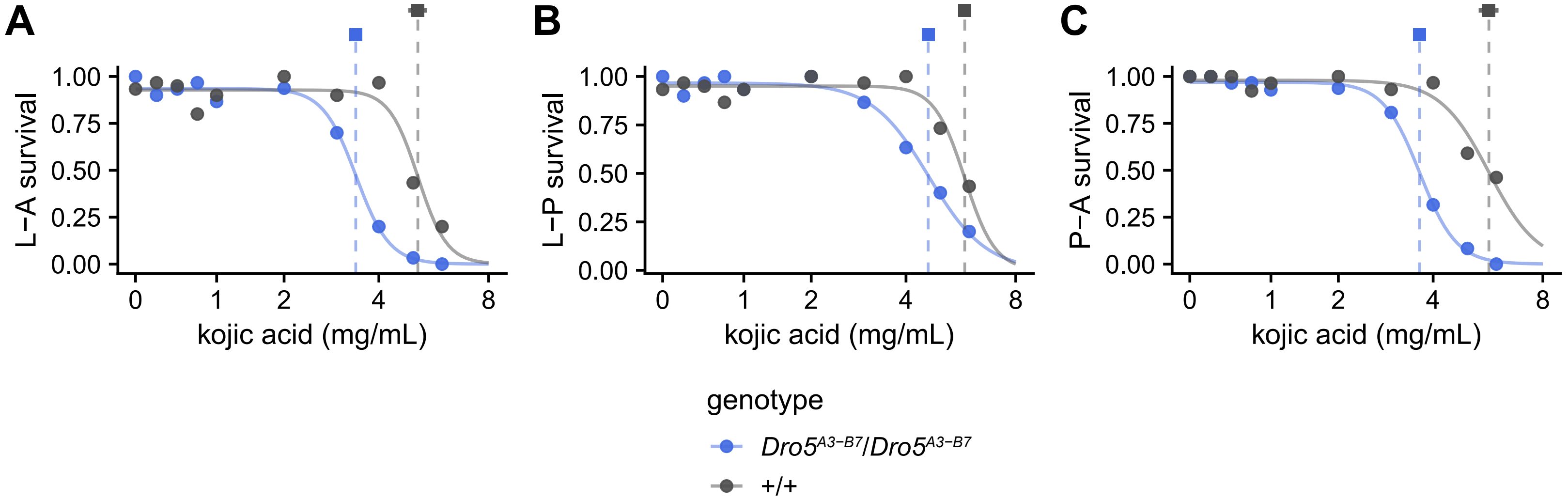

### Figure 7

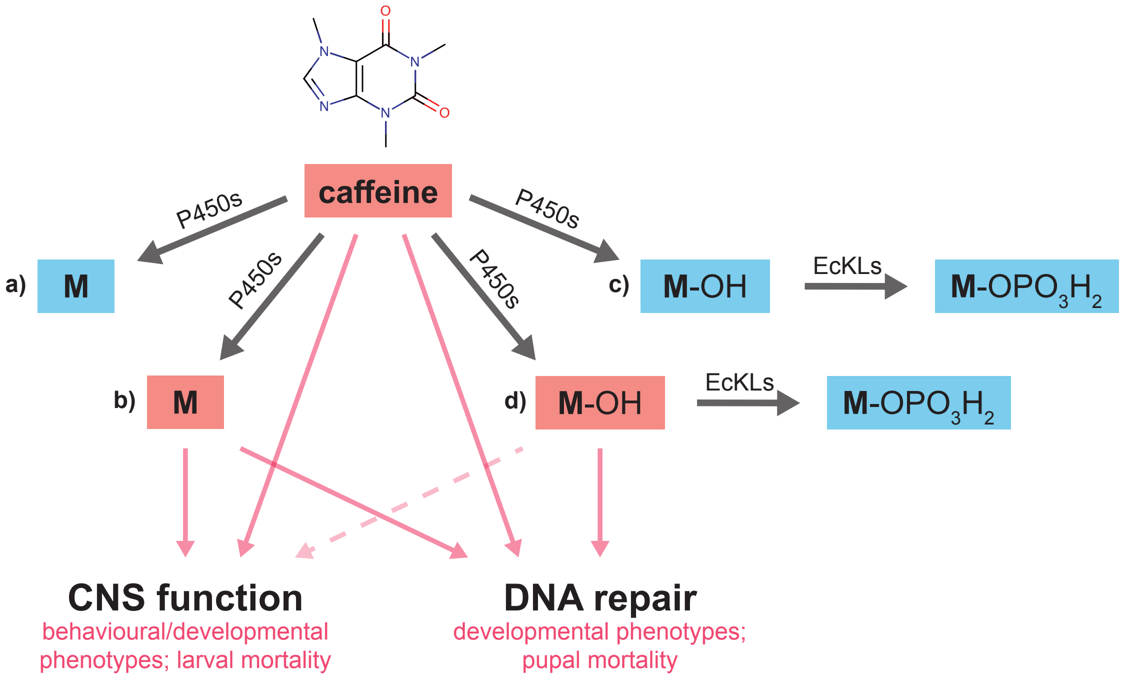

### Supplementary Figure 1

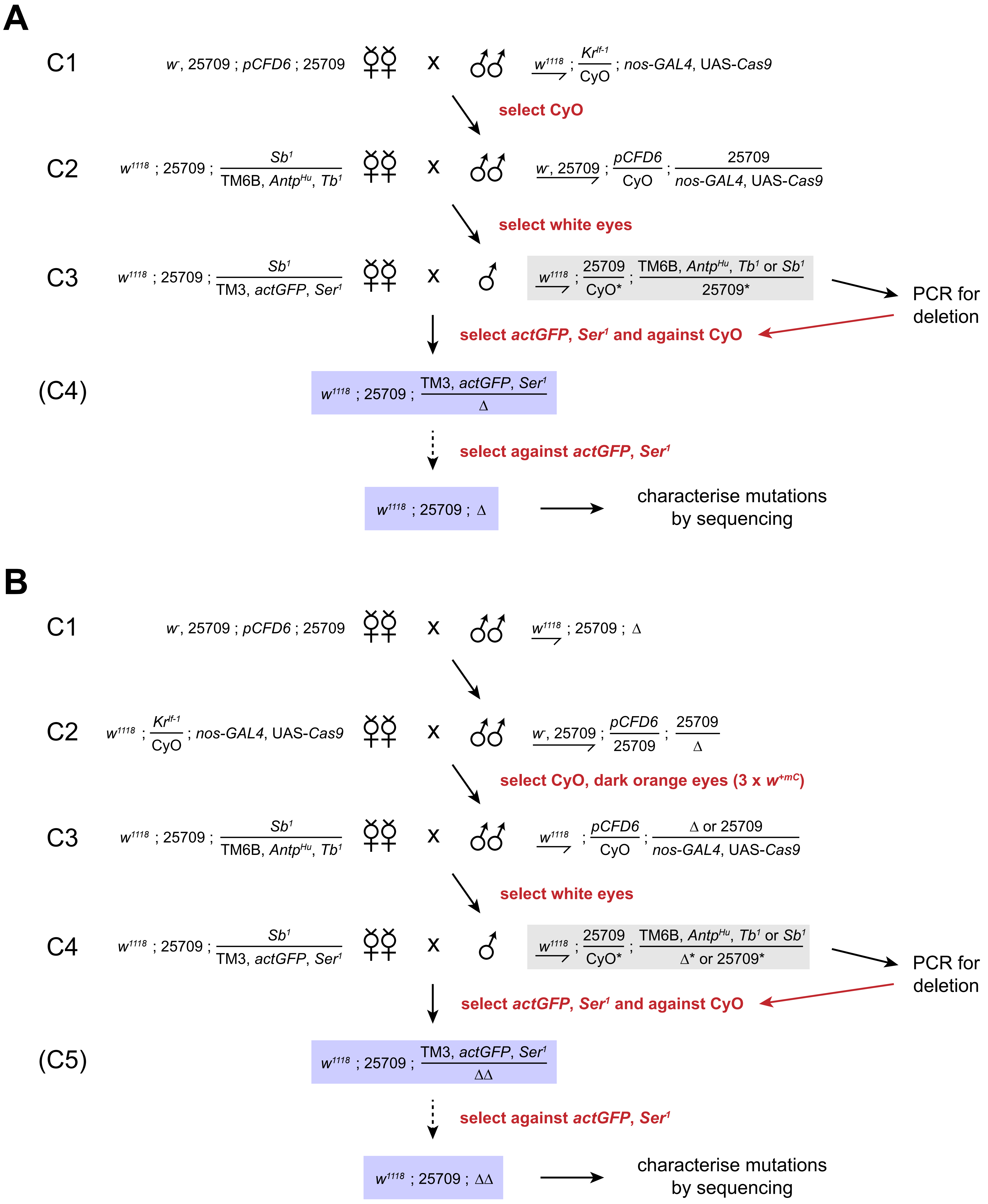

### Supplementary Figure 2

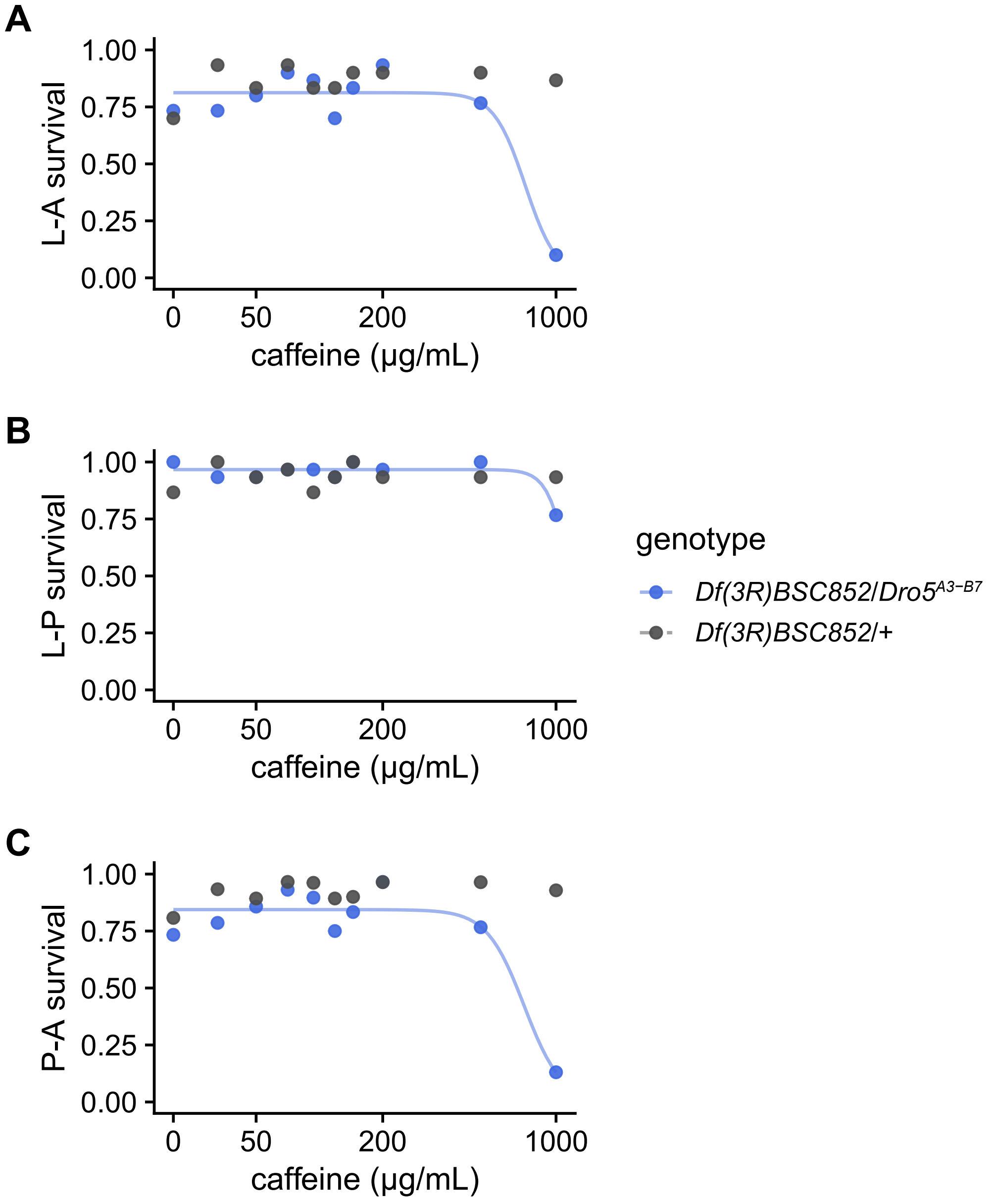

### Supplementary Figure 3

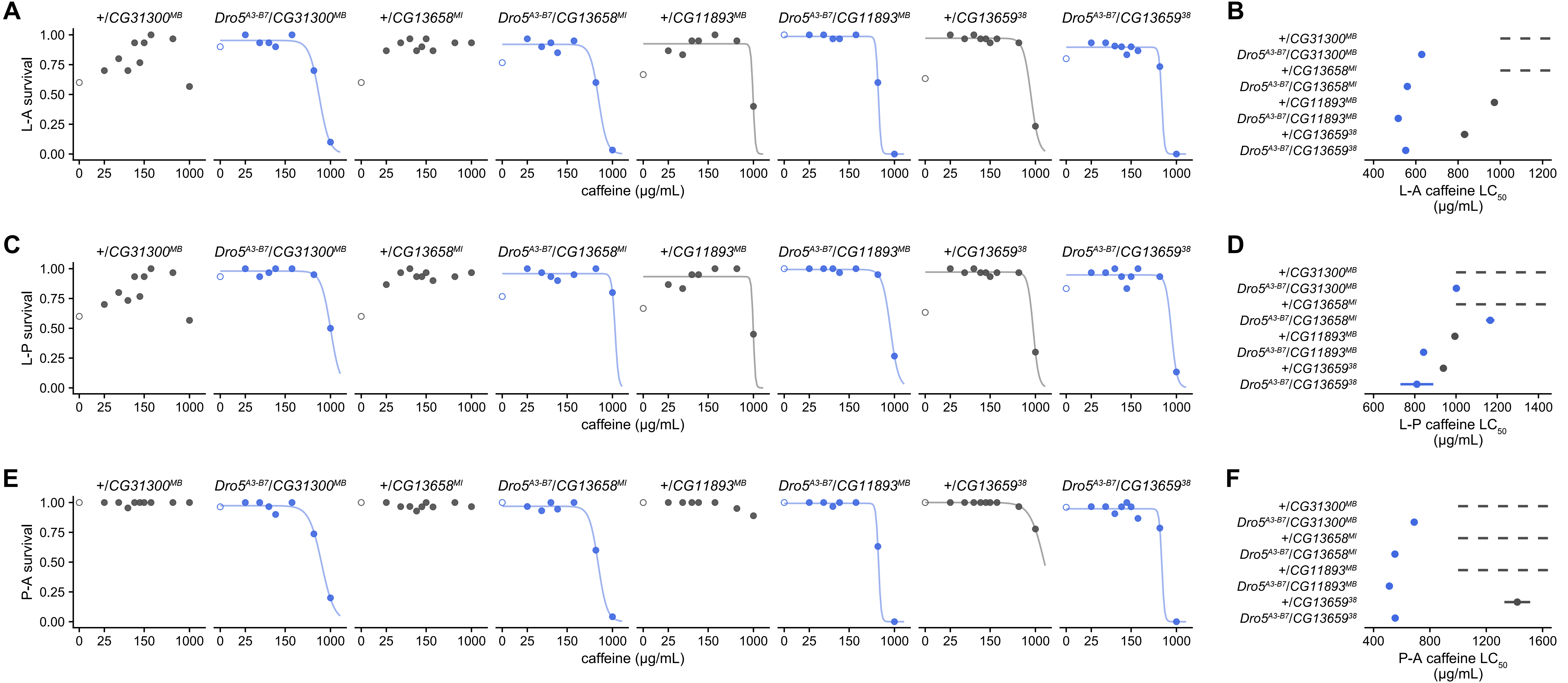

### Supplementary Figure 4

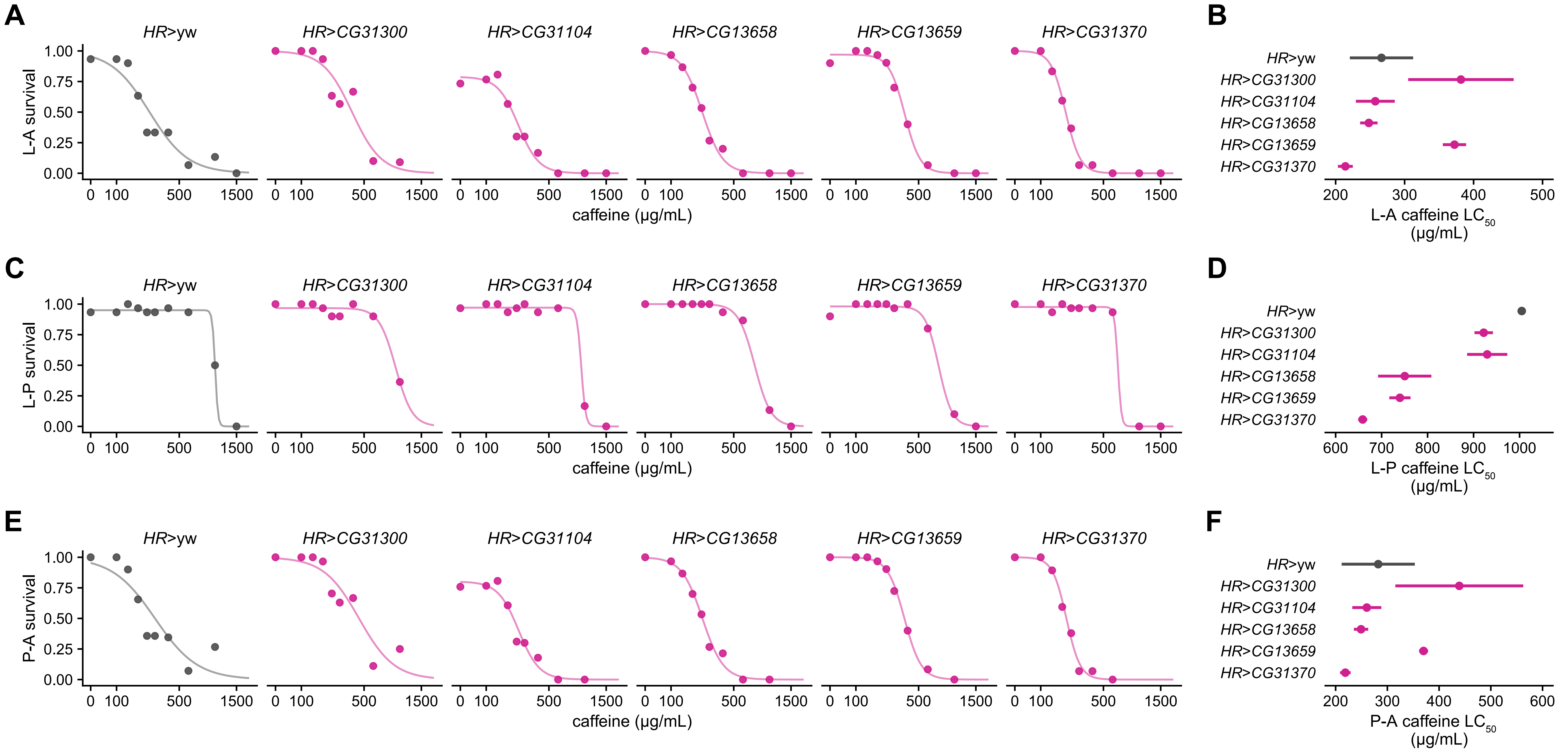

### Supplementary Figure 5

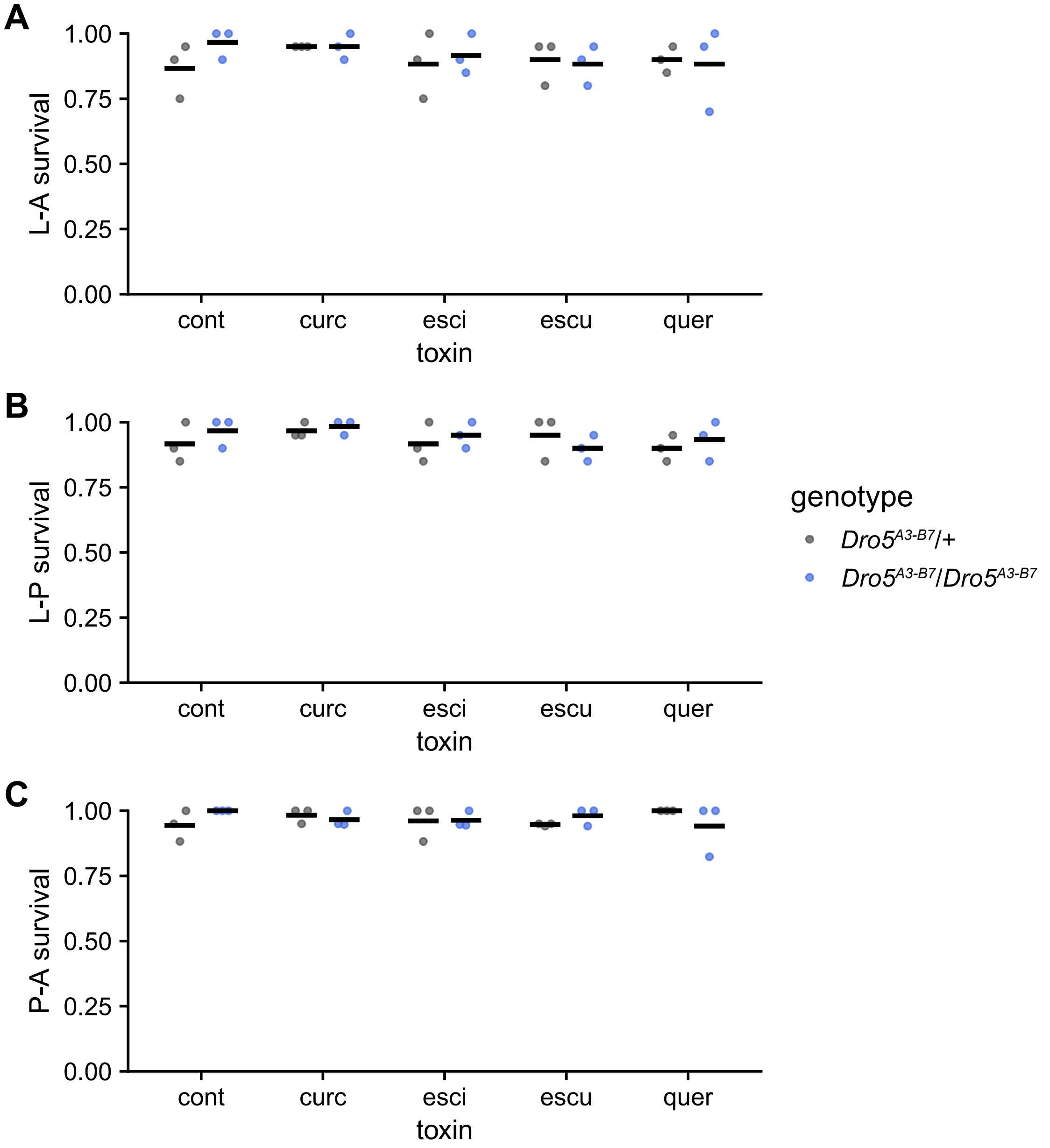

### Supplementary Figure 6

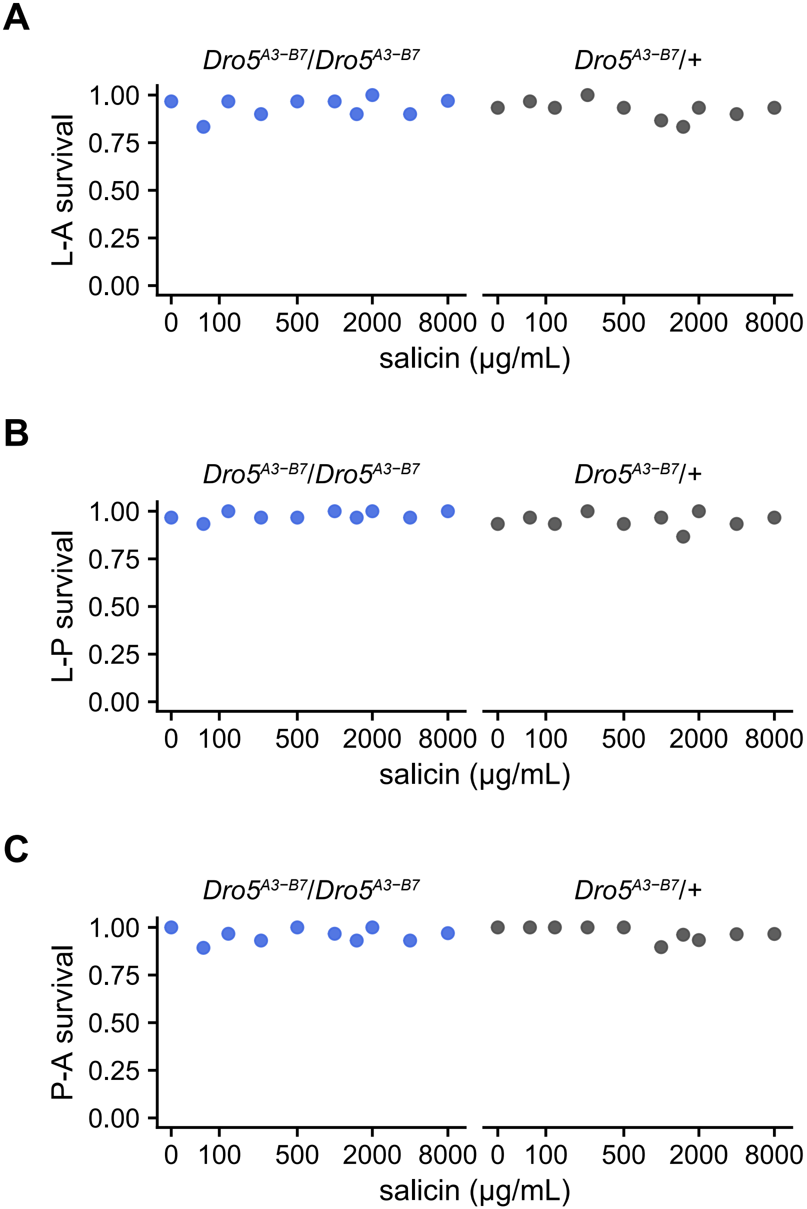

### Supplementary Figure 7

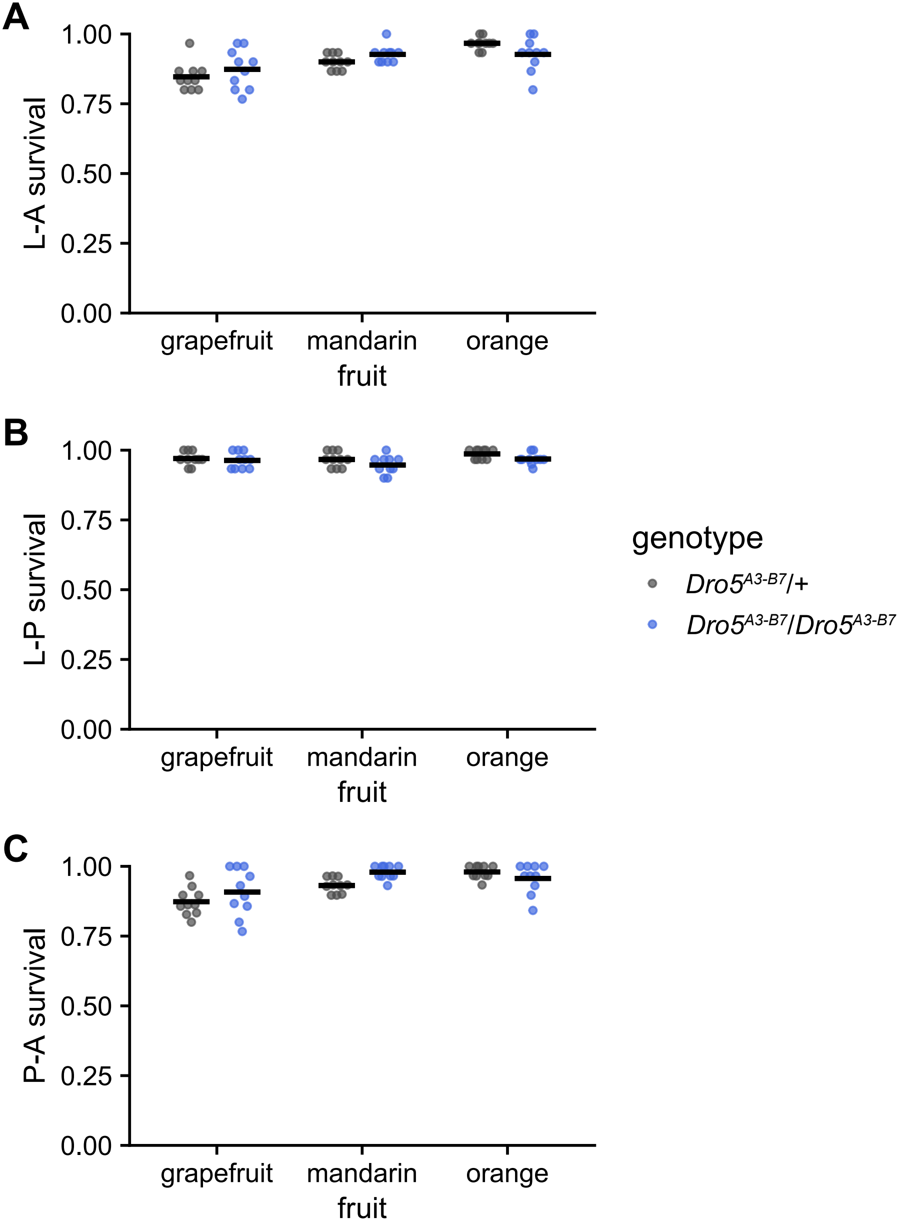
